# A systems-based approach to uterine fibroids identifies differential splicing associated with abnormal uterine bleeding

**DOI:** 10.1101/2024.02.06.578872

**Authors:** CY Wang, M Philpott, DP O’Brien, A Ndungu, J Malzahn, M Maritati, N Mehta, V Gamble, B Martinez-Burgo, S Bonham, R Fischer, K Garbutt, CM Becker, S Manek, AL Harris, F Sacher, M Obendorf, N Schmidt, J Mueller, T Zollner, KT Zondervan, BM Kessler, U Oppermann, AP Cribbs

**Author notes:** Equal contribution (CY Wang, M Philpott).

## Abstract

Uterine fibroids (UFs), benign tumours prevalent in up to 80% of women of reproductive age, are associated with significant morbidity, including abnormal uterine bleeding, pain and infertility. Despite identification of key genomic alterations in MED12 and HMGA2, the pathogenic mechanisms underlying UFs and heavy menstrual bleeding (HMB) remain poorly understood. To correlate systematically genetic, transcriptional and proteomic phenotypes, our study involved an integrative analysis of fibroid, myometrium and endometrium tissues from 137 patients, utilising genome-wide SNP arrays, targeted sequencing, RNA sequencing and proteomics. Our findings reveal 39.7% of UFs possess MED12 mutations, alongside novel variants in genes such as COL4A5 and COL4A6. Multi-omics factor analysis of integrated protein and mRNA highlighted differential regulation related to extracellular matrix remodelling, proteolysis and homeostasis in fibroid versus myometrium tissues, and distinct gene sets associated with RNA splicing in the endometrium of patients with HMB, particularly in MED12-mutated fibroids. Our study proposes a model, which is supported by *in vivo* evidence, where altered signalling of MED12-mutated fibroids influences RNA transcript isoform expression in endometrium, potentially leading to abnormal uterine bleeding. This integrative approach unravels complex molecular pathways in UF pathogenesis and HMB, offering novel insights for targeted therapeutic development.

## INTRODUCTION

Human uterine fibroids (UF), also known as uterine leiomyoma, are benign tumours of the uterus that affect a large population of women of reproductive age. They are particularly prevalent in Black women in the United States, with an incidence of approximately 80% for those aged between 35 and 49, which is significantly higher than the rate of 70% for white women of the same age ^1^. UFs interfere with normal uterine function, and in more than half of cases can cause distressing symptoms such as heavy menstrual bleeding (HMB), pelvic pain, urinary incontinence, and/or infertility ^2^. Despite the high prevalence of the condition, treatment options are hindered by the broad range of clinical manifestations. Symptomatic UFs are removed either by hysteroscopic/laparoscopic surgery, embolization, or hysterectomy ^3^. In the United States alone, UFs are cited to be the cause of over 50% of hysterectomies ^4^, and direct costs for their treatment have been estimated to be between $4-9 billion annually ^5^. Irregular heavy menstrual bleeding (HMB; or AUB, abnormal uterine bleeding) is the most common symptom of UFs (46%; ^6^) and adversely affects quality of life as a result of concurrent pain, anaemia, mood swings, and potential social embarrassment ^7–9^. Despite the apparent link between Ufs and HMB, the pathomechanisms behind the two remain unclear ^9^.

Mutually exclusive driver mutations in the mediator subunit 12 (MED12) ^10^ and high-mobility group AT-hook 2 (HMGA2) ^11^ genes have been identified in human Ufs that, when combined, occur in ∼90% of disease cases. MED12 forms part of the Mediator Complex, which regulates transcription initiation and elongation by RNA polymerase II ^12^, while HMGA2 binds to, and alters the structure of DNA, promoting assembly of protein complexes that regulate transcription ^13^. Additional genes have been implicated in UF, e.g. genetic inactivation of fumarate hydratase (FH), a key enzyme of the Krebs cycle which can impair cellular metabolism and promote hypoxia, imparts a higher risk of Ufs ^14,15^. Furthermore, deletion of the collagen genes COL4A5 and COL4A6 are associated with a familial UF aetiology ^11,16^. The underlying causes of how mutations in certain genes lead to the development of Ufs and associated symptoms, are not yet fully understood. Accordingly, gaining a deeper understanding of these mechanisms could potentially reveal key targets for therapeutic intervention and provide opportunities for biomarker identification.

To address the gap in understanding the underlying mechanisms of UFs, a number of studies have applied “omics” techniques to investigate the mechanisms of the condition. These studies have primarily used microarray technology to compare gene expression profiles between myometrium and UF, but the sample size of these early studies was often limited^17–24^. Recent studies have built upon the growing knowledge of UF genotypes to gain deeper insights into the condition. A study by Mehine et al. (2016), for example, explored the transcriptional differences in UF from 60 patients carrying different genetic drivers, such as MED12 mutations, HMGA2 rearrangements, FH inactivation and collagen gene deletions. This research revealed specific changes in key pathways, including Wnt, prolactin, and insulin-like growth factor 1 (IGF-1) signalling, that were unique to different subsets of UFs. While a number of studies have used proteomic approaches to compare normal myometrium and UFs, these efforts have been hindered by small sample sizes, with patient cohorts ranging from as few as 3 up to 10 patients ^25–31^. Despite this, these studies have provided valuable insights into the role of processes such as apoptosis, inflammation, and cytokine regulation in the development of UFs. Taken together, the majority of “omics” studies have pointed to a link between the development of UF and the biology of extracellular matrix (ECM), the signalling pathways of growth factor β3 (TGF-β3) and WNT-β-catenin^28–31^.

To further understand the mechanisms of UFs and HMB, we first set out to significantly increase sample size and included matched samples for endometrium, myometrium and fibroid using tissue from 94 UF and 43 non-UF patients. By utilising data integration tools, we were able to relate transcriptomic, proteomic, genotype and clinical information to provide further insights into the pathomechanisms of UFs.

## RESULTS

### Clinical features of the cohort

A total of 137 patients undergoing hysterectomy, myomectomy or trans-cervical resection of fibroids (TCRF) were recruited into the study (Fig. 1a). The majority of these patients underwent surgery for the removal of UFs, but material was also collected from 43 non-UF patients (Fig. 1b). These patients serve as a comparative non-UF cohort for certain analyses. However, its crucial to note that they did not represent healthy controls; instead, they underwent hysterectomies for a variety of conditions including endometriosis, adenomyosis, ovarian cysts or cervical neoplasia (Supplementary Table 1). HMB status was determined from a combination of patient questionnaires and clinical notes, with HMB established for 95 donors (Fig. 1c). For subsequent analyses, it was important to distinguish between historical HMB (often absent at the time of surgery due to hormonal treatments, but still relevant to genetic analyses) and HMB during the menstrual cycle in which surgery took place (relevant to phenotypic analyses). Menstrual cycle phase for patients was primarily determined by histological examination of endometrium, but where this was not possible, clinical notes, progesterone/estradiol ratios from blood samples or self-reported date of last menstruation was used (Fig. 1d). A high proportion of the donors had inactive endometrium (i.e., no active menstrual cycle) due to hormone treatment for UFs or associated HMB. Material collected from surgeries was macroscopically prepared into endometrium, myometrium, UF and the pseudocapsule region between the myometrium and fibroid, although not all tissue types could be collected from all samples. Samples collected by TCRF tended to be of poor quality and yielded little or no endometrium, as did myomectomies, and surgeries performed by morcellation could not be reliably separated into individual tissue types. Overall, tissues from 91 donors were retained for this study and deemed suitable for multi-omics analysis (Table 1).

**Fig. 1:**
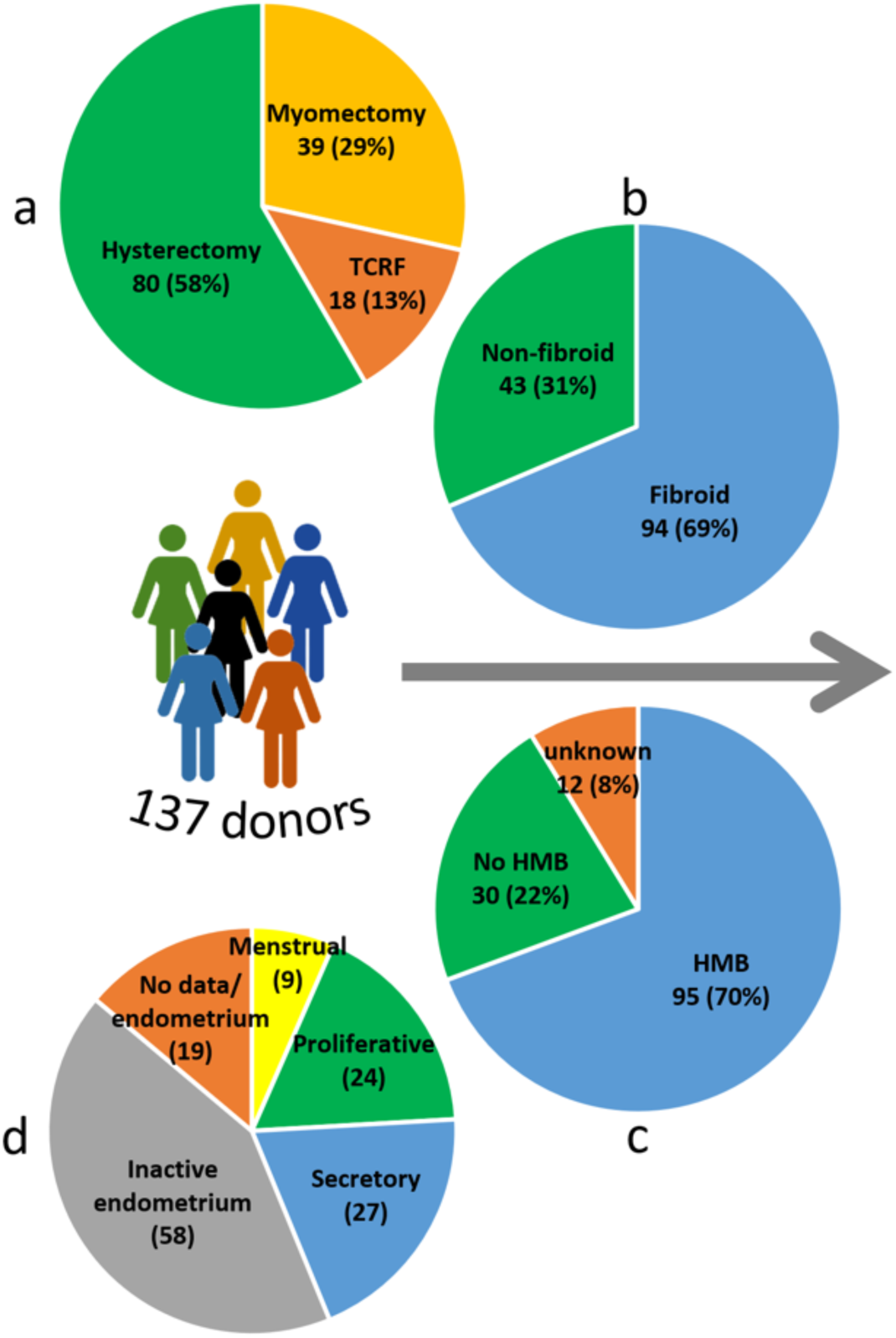
Tissues collection used in this study. Breakdown by **a)** type of surgery, **b)** fibroid vs control (non-fibroid) patients, **c)** HMB status and **d)** menstrual cycle phase.

**Table 1:**
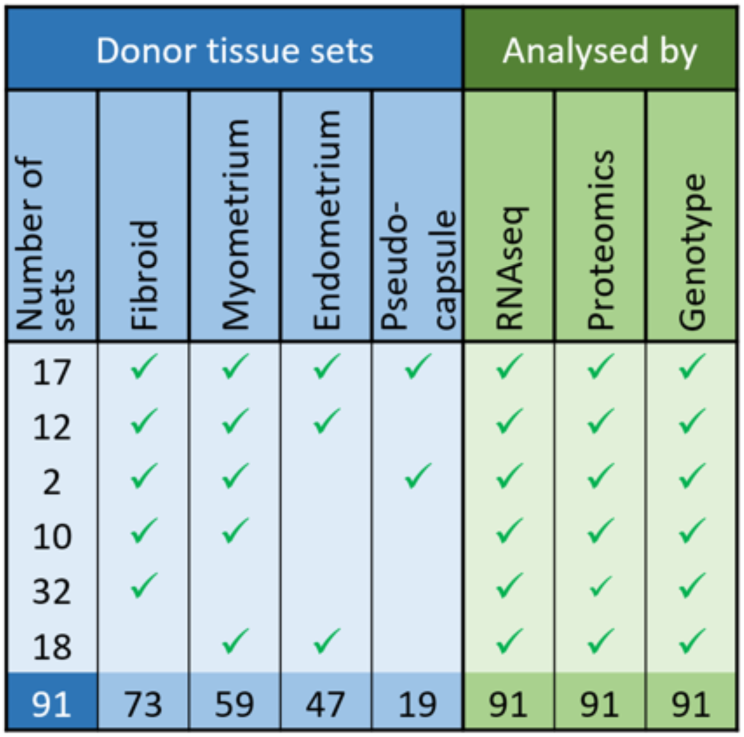
Tissues form 91 donors were suitable for multi-omics systems analysis and the table depicts the tissue sets collected and the analyses performed on the tissues.

### Genomic analysis suggests multiple lesions and novel candidates and pathways in UF pathology

We deployed a targeted sequencing approach for genes which have been reported to be involved in UF formation, with the target panel spanning MED12, HMGA2, FH, COL4A5/6, HMGA1^32^, RAD51B^11^, AHR^33^, CAPRIN1^33^, CUX1^34^, DCN^33^ and PCOLCE^35^ candidate genes (Supplementary Table 2) . The analysis focused on single nucleotide polymorphism (SNPs), point mutations and short indels within those gene loci (Fig. 2a-b, and Supplementary Table 3). Among 73 fibroids, 29 (39.7%) were identified with MED12 common UF mutations located at intron 1 and exon 2. Other mutations in the MED12 coding regions were either synonymous or situated at splice regions with minimal impact. Our comprehensive analysis unveiled a total of 55 distinct mutations within the coding regions of MED12 (Fig. 2a). In addition to MED12, we identified variants predicted to impact protein function across our target panel, each with various frequencies (Fig. 2a and Supplementary Fig. 1b). Notably, specific hotspots were observed in COL4A6 and CUX1 (Fig. 2b). Within exon 24 of COL4A6 (chrX: 108183752, 108183753 and 108183754), 67 fibroids exhibited in-frame insertion, deletion-insertion and/or frameshift variants. Notably, at the locus of chrX: 108183753, 45 displayed heterozygosity wherein 39 showing a wildtype/ in-frame deletion-insertion mutation (where Glu657 was replaced by two amino acid residues Asp and Lys), and 6 carrying this in-frame deletion-insertion along with the frameshift mutation after Val658. Furthermore, 2 fibroids displayed homozygosity of the in-frame deletion-insertion at Glu657. Another hotspot in COL4A6, encoding a missense variant (Ser > Pro), was identified in exon 20 (Fig. 2b). In conjunction with these findings in COL4A6, our analysis revealed 60 fibroid samples harbouring missense variants in CUX1, particularly at exon 16 (Fig. 2b, bottom panel).

**Fig. 2:**
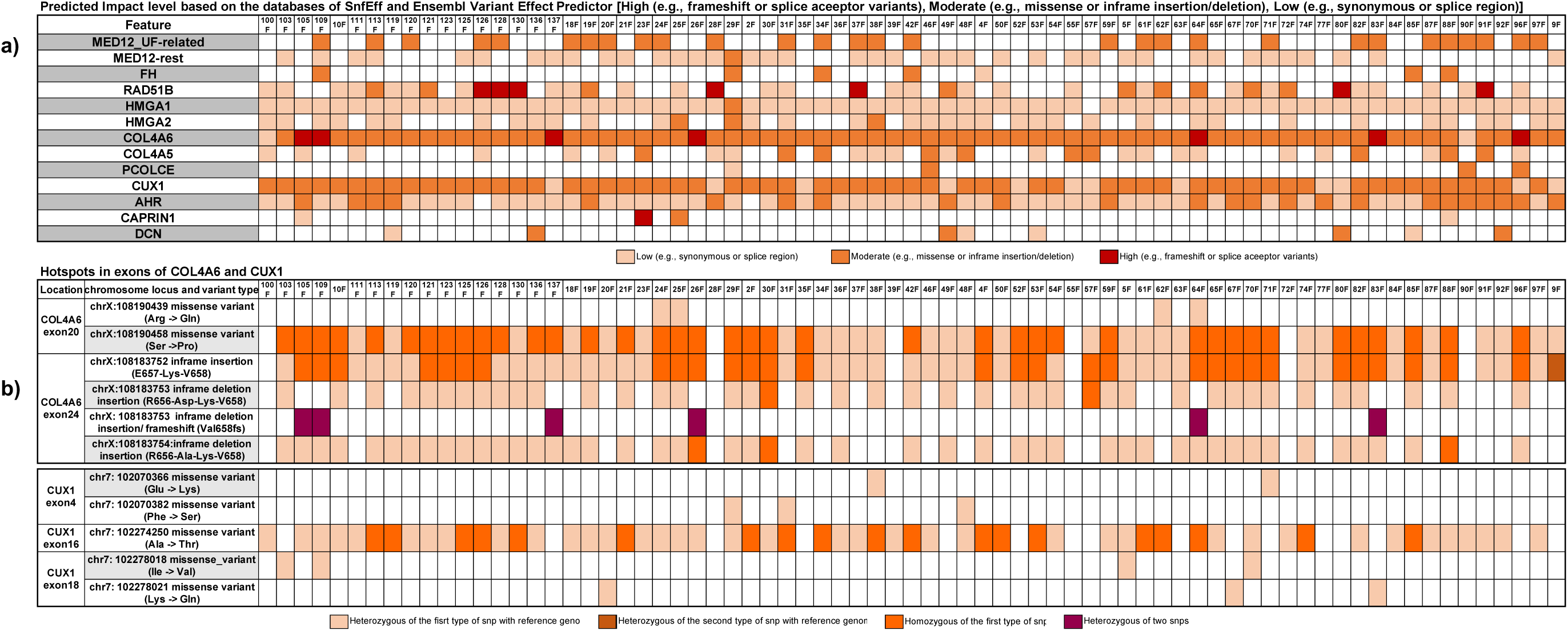
Fibroid genotyping by targeted sequencing. **a** Mutations of known fibroid-related genes in the fibroid samples were identified by SureSelect targeted sequencing. Impact level prediction of identified variants on protein-coding is marked in high (colour in red), moderate (colour in orange) and low (colour in light orange). **b** Hot spots of variants detected in COL4A6 and CUX1.

We further investigated variants which, despite being predicted to exert minimal impact on protein functionality, exhibited correlations with distinct phenotypes. These included fumarase deficiency, metabolism, hormone, cancer deposition and leiomyomatosis (Supplementary Fig. 1a,c). These phenotype associations were drawn from diverse sources including ClinVar^36^, dbVar^37^, and the NHGRI-EBI GWAS catalogue ^38^, all integrated into the Ensembl Variant Effect Predictor^39^. In our analysis of COL4A6, we identified several homozygous SNPs associated with X-linked Alport syndrome-diffuse leiomyomatosis (AS-DL)^40^ at the loci (ChrX: 108173940, 108204943, 108247572 and 108259167). Additionally, a limited number of fibroid samples exhibited similar genetic association with AS-DL in COL4A5, as shown in Supplementary Fig 1a, c and Supplementary Table 4. Exploring structural variants within our targeting sequencing dataset revealed the presence of various deletions ranging from 10 bp to several hundred bps, with no identified translocation events (Supplementary Fig. 1d). Collectively, our targeted sequencing analysis unveiled a distinctive distribution of driver mutations within our cohort, showcasing an unexpected scarcity of aberrations in MED12, and a notably higher occurrence of collagen variants.

Complementing our targeted sequencing approach, we utilised short-read RNA-seq data to broaden our investigation into potential chromosomal translocations within the UF samples. Employing Arriba^41^ for this analysis, we revealed several fusion events identified with medium to high confidence in 21 fibroids, as detailed in Supplementary Table 5. HMGA2 translocation was identified in one fibroid sample (Table 2) and was subsequently validated through long-read whole-genome sequencing.

**Table 2:**
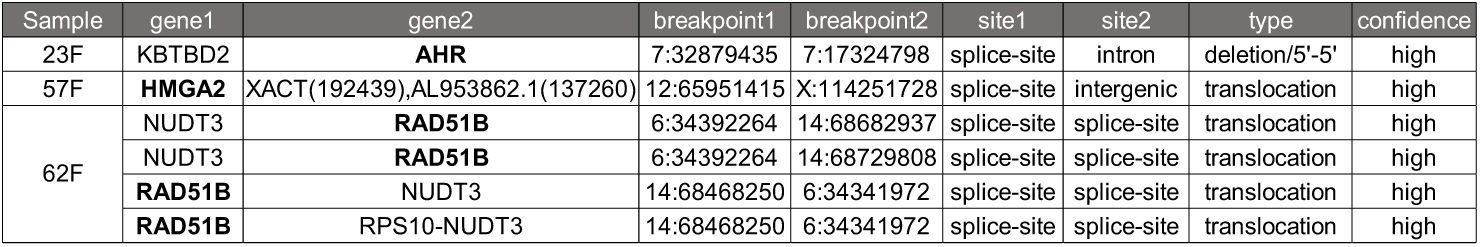
Predicted translocation and mutations in the fibroid samples using bulk RNA-seq data.

### Omics analysis of UF myometrium and fibroid tissues highlights changes in angiogenesis, ECM and signalling pathways in UF patients

Bulk short-read RNA-seq analysis was performed on 165 samples, encompasing 63 fibroid, 57 myometrium, and 45 endometrium tissues. Post stringent quality control, a subset of 120 samples (Supplementary Table 7) were selected for in-depth transcriptomic analysis. Comparative transcriptomic profiling between UF myometrium and fibroids (Supplementary Fig. 2) employed gene set enrichment analysis (GSEA) and gene ontology (GO), pinpointing significant activity in processes such as angiogenesis and blood vessel endothelial cell migration (Supplementary Fig. 2b). Additionally, over-representation analysis of 600 genes exhibiting differential expression (padj < 0.05, absolute log2 fold change (abs(log_2_FC)) ≥ 1.5) underscored pathways related to ECM-related pathways, hormone metabolic process, and regulation of signalling receptor activity (Supplementary Fig. 2c). At the proteomic level, applying a 1% false discovery rate (FDR), we identified 310 proteins as differential expressed in fibroid versus UF myometrium (Supplementary Fig. 3a). The pathway delineation of these proteins revealed associations with ECM organisation, collagen fibril biology, blood vessel morphogenesis, platelet degranulation, VEGF and ErbB signalling as well as oxidative stress (Supplementary Fig. 3b).

Next, we aimed to better understand the impact of MED12 UF mutations on the pathology of fibroid (Supplementary Fig. 4). In the over-representation analysis of 97 differentially expressed genes (padj < 0.05, abs(log_2_FC) ≥ 1.5) derived from bulk transcriptomics, we identified significant associations within the regulation of systemic arterial blood pressure, implicating MED12 UF mutations potentially influence or alter blood vessel function (Supplementary Fig. 4b).

### Endometrial mRNA expression profiles can be distinguished based on HMB status and highlight enrichment of immune processes during secretory phase

Principal component analysis (PCA) of bulk transcriptomics (Supplementary Fig. 5a) exhibited a distinct separation between UF patients experiencing HMB with active menstrual cycle (n=15) and those without HMB along the PC1 and PC2 axes. We identified 175 differentially expressed genes (padj < 0.05, abs(log2FC) ≥ 1.5) in this context. GSEA highlighted top-enriched pathways, notably vasculature development, blood vessel morphogenesis and extracellular structure organisation (Supplementary Fig. 5b). Similarly, the HMB status of proliferative endometrial tissues from both UF and non-UF patients was compared using the proteomic dataset, revealing 135 differentially expressed proteins (Supplementary Fig. 6a). Pathway analysis indicated the up-regulation of translation/RNA/mRNA metabolic processes, protein folding and turnover, muscle contraction and oxidative phosphorylation (Supplementary Fig. 6b). Conversely, several pathways were downregulated in HMB proliferative endometrial tissues compared to corresponding non-HMB samples. These included processes related to cytokine production, angiogenesis, blood clotting, as well as ECM degradation. Next, we analysed the impact of HMB on endometrial mRNA expression profiles stratified by menstrual cycle phase (Supplementary Fig. 7-8). Whilst changes in cell cycle and mitosis are dominant during the proliferative phase (Supplementary Fig. 7), GSEA highlights during secretory phase (Supplementary Fig. 8) immune processes such as inflammatory response or allograft rejection besides RAS signalling as significantly enriched when HMB vs non-HMB status are compared. Inflammatory processes and leukocyte trafficking in the endometrium during the menstrual cycle^42^, are supported by our finding which implicates contribution of immune responses on HMB, highlighted by IL-11 for example, which was differentially expressed in HMB particularly in the secretory phase (Supplementary Fig. 8c).

### Multi-omic factor analyses identifies RNA processing and confirms ECM as major factor in fibroid biology

Utilising multi-omic factor analysis (MOFA)^43,44^, we integrated bulk transcriptomics and proteomics data, enabling enhanced analytical depth within individual datasets. This approach facilitated nuanced comparisons between fibroid and myometrium tissues from patients, irrespective of UF presence, and extended to include endometrial tissues from both UF-affected and unaffected patients.

Upon conducting an integrative analysis of fibroid (n=50) and myometrium (n=40, 30 with UF and 10 without), 8 factors were defined based on observed variances among the samples (Fig. 3a). Despite utilizing batch corrected data as input, a batch effect was detected in three factors (factor 1, 3 and 4), either in transcriptomics or proteomics, underscoring the sensitivity of MOFA analysis to hidden underlying factors. Factor 2, which is devoid of batch effects (Supplementary Fig. 9b), exhibited a robust correlation with tissue type (myometrium or fibroid) (Fig. 3b-c). GSEA reveals enriched pathways, consistently implicating ECM organisation, vasculature development, in addition to WNT signalling, wound healing, and immune response (Supplementary Fig. 9a), aligning with our previous analyses. A set of 52 features further highlighted ECM organisation, angiotensin maturation, and hormone metabolism (Fig. 3d-e). Additionally, among the features positively associated with fibroid, a predominant number were components of the ECM, including structural proteoglycans like versican (VCAN) and collagen proteins (Supplementary Fig. 10). These findings were reinforced through another integrative analysis conducted using 2D Annotation Enrichment analysis^45^ integrated in the Perseus software for proteomic analyses, yielding congruent results. (Supplementary Fig. 11).

**Figure 3.**
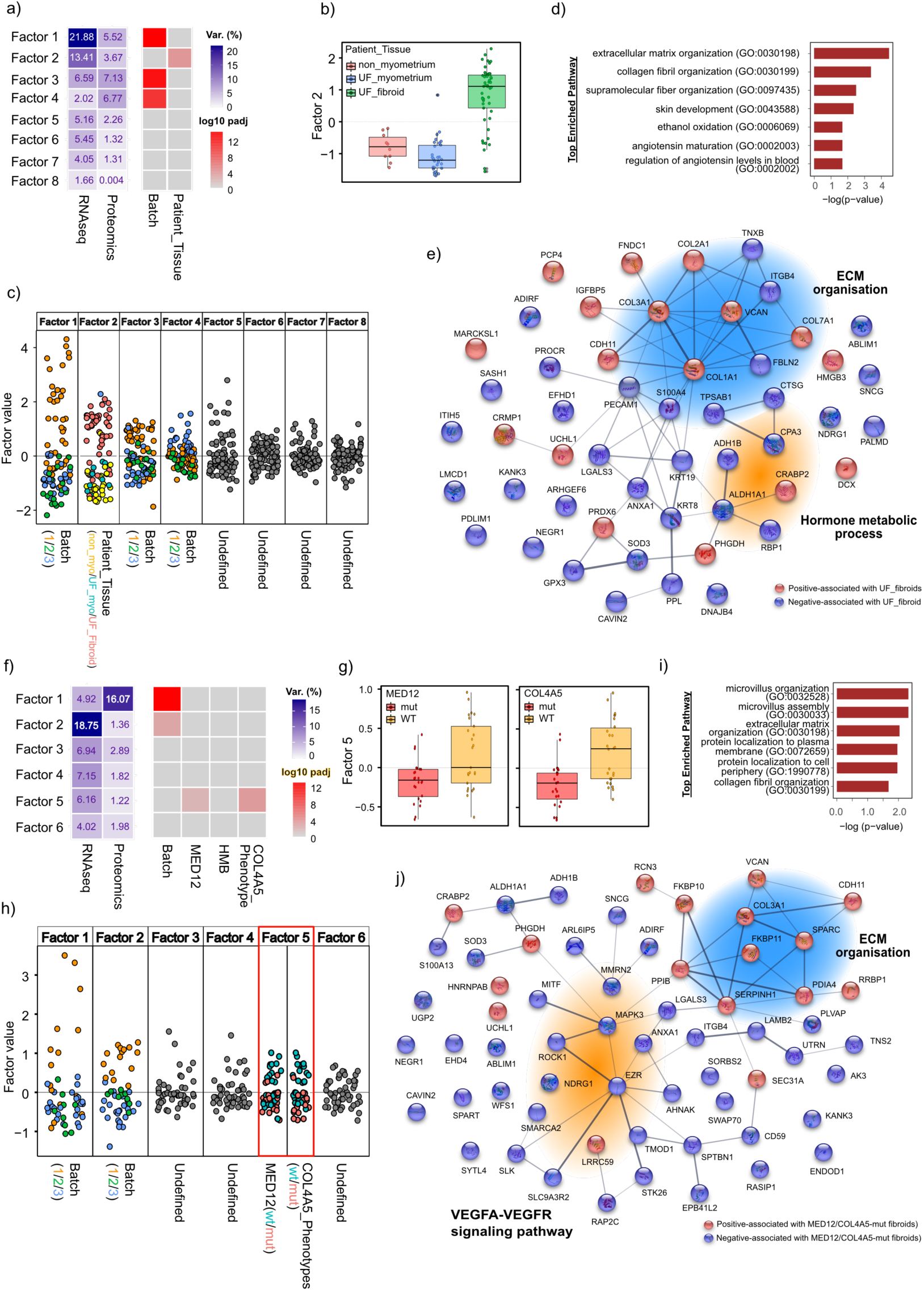
Systems level analysis of myometrium vs fibroid. (top; a-e) and fibroids (mutants) (bottom; f-j) Relative contributions of transcriptomic and proteomic datasets to MOFA factors (% variance) **a, f left)**. Correlation of variance to indicated parameters (Log_10_ padj) **a, f right)**. Boxplot of the sample groups in the indicated factor **(b, g)**. Scatter plot of factor values differentiated by indicated parameters. **c, h)**. Top enriched pathways using MSigDB C5 GO:BP **(d, i)** and STRING diagram **(e, j)** depicting relationships between top-weight genes from systems analysis.

To gain deeper insights into the factors underlying fibroid formation, we refined MOFA analysis specifically within fibroid tissue. Among the 6 factors defined by this analysis, factor 5 exhibited a noteworthy albeit weak correlation with the mutational status in MED12 (MED12 UF mutations, n=24; MED12 WT, n=25) and COL4A5 phenotype (SNPs linked to AS-DL, n=23, and WT (the reference genome), n=26) (Fig. 3f-h and Supplementary Fig. 12b), akin to observations in the transcriptomics PCA plot (Supplementary Fig. 4a and 13). Consistent with the results from GSEA (Supplementary Fig. 12a), the enriched pathways of 61 features highly associated with mutant fibroid highlighted ECM organisation and collagen fibril organisation. This finding implies that fibroids harbouring either MED12 UF mutation or COL4A5 SNPs related to AS-DL, compared to WT (i.e. non mutated in MED12 or COL genes) fibroids, potentially have a higher impact on dysregulation of genes involved in ECM deposition, collagen biosynthesis and collagen chain trimerization, such as SERPINH1 (Serpin Family H Member 1) and PPIB (Peptidylprolyl Isomerase B), respectively (Supplementary Fig. 14).

To better understand the occurrence of HMB in UF patients, we conducted MOFA analysis encompassing all endometrial samples from patients with and without UF (n=31). Among the 7 factors (Fig. 4 & Supplementary Fig. 15b), factor 1 demonstrated a correlation with HMB and hormone treatment history, factor 2 exhibits a correlation with the presence of fibroid, highlighting the influence of fibroid tissue on physiological functions of endometrium, while factor 7 correlates not only with the presence of fibroid but also stratifies fibroids based on their MED12 mutational status, further correlating with HMB (Fig. 4a right & Fig. 4b-c). The finding on factor 1 suggests the potential impact of therapeutic interventions on HMB or their influence on endometrium function; the effect lasts even after treatment cessation. Features highly associated with factor 1 participated in angiogenesis, wound healing, and ECM organisation (Fig. 4d, top panel); a majority displayed a negative association with HMB (Fig. 4e and Supplementary Fig. 16). For example, CD59, whose genetic deficiency results in hemolytic anemia and thrombosis^46^, and angiogenin (ANG), a member of the RNAase A superfamily, has the ability to induce neovascularization^47^.

**Fig 4:**
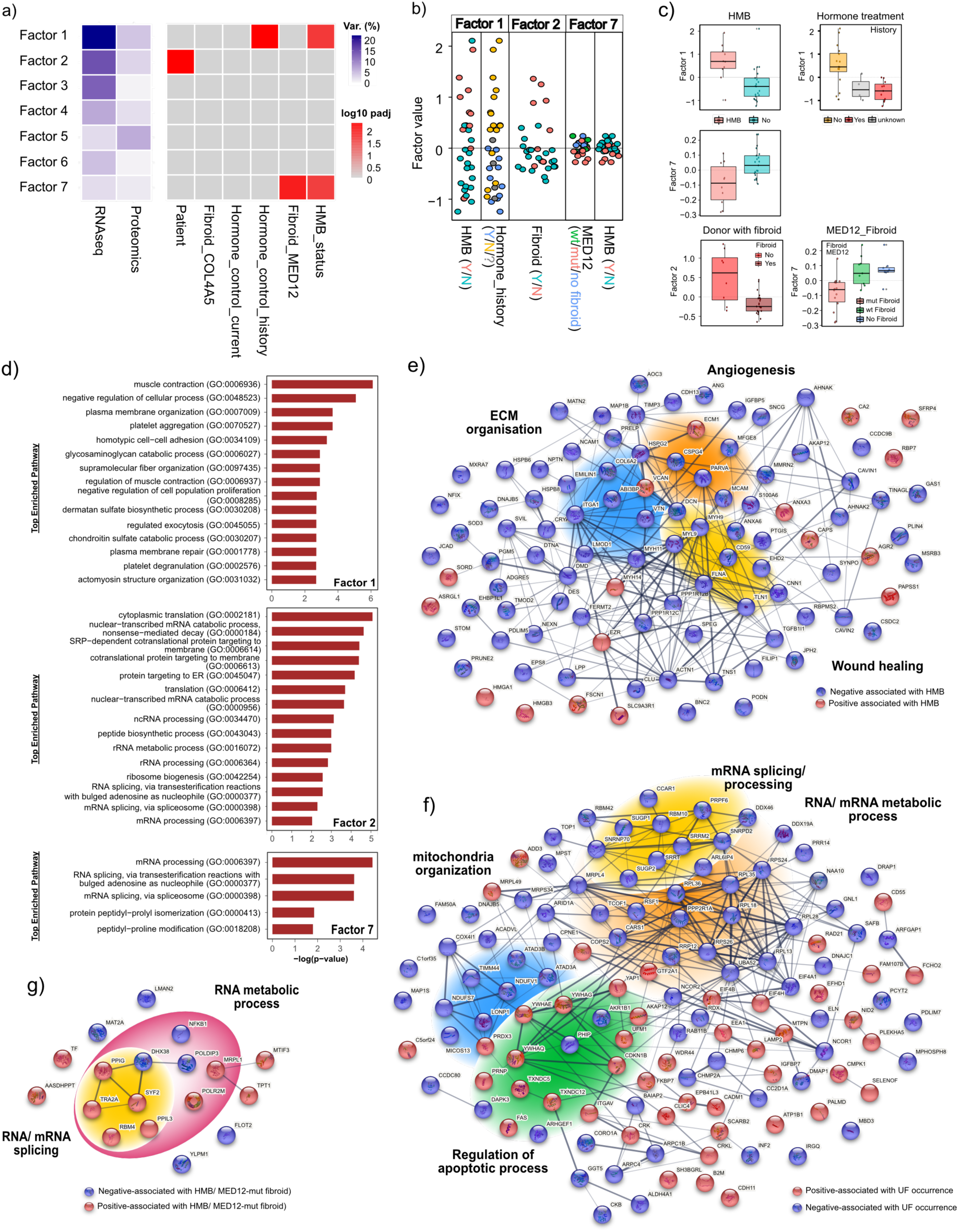
Systems level analysis of endometrium (proliferative, secretory phase and inactive endometrium from UF and non-UF patients) **a)** Relative contributions of transcriptomic and proteomic datasets to MOFA factors (% variance; left panel) and correlation of variance to indicated parameters (Log_10_padj; right panel). **b)** Scatter plot of factor values differentiated by indicated parameters. **c)** Boxplot of the sample groups in the indicated factor. **d)** pathway analysis **e-g)** STRING diagrams depicting relationships between the features associated with the correlated parameter.

Interestingly, pathway analysis revealed that features in factor 2 and factor 7 were predominantly involved in RNA processing and metabolic process, including mRNA splicing and RNA 3’-end processing (Fig. 4d, middle and bottom panel, 4f-g and Supplementary Fig. 17-19). In summary, our multi-omics integrative analysis indicates a potential disruption of RNA homeostasis and the subsequent splicing events in endometrium due to UF occurrence. This disruption is further exacerbated by MED12 UF mutations, potentially enhancing these disturbances.

### Differential transcript usage in the endometrium obtained from patients carrying MED12-UF mutant fibroids

Our integrative analysis identified the involvement of RNA processing and mRNA splicing in UF endometrium pathology. To scrutinize transcript-level alterations in UF endometrium with active menstrual cycle (n=15), excluding therapeutic intervention, we conducted differential transcript usage (DTU) analysis. Comparing samples from patients with MED12-UF mut fibroid and those with no mutations in MED12, we observed differential expression of transcript variants in splicing-related genes (RBM4, PPIG and HNRNPC) (Fig. 5a). The DTU analysis identified 2,111 genes, with top enriched pathways as RNA processing/splicing, mRNA transport, and TGF-β receptor signalling pathway (Fig. 5b). We identify altered transcript usage in genes like ANGPT1, ANGPT2, TGFBR1, TGFBR2 and TGFBR3 in the endometrium from patients with MED12-UF mut fibroid (Fig. 5c-d), emphasizing the impact on fibroid mechanisms due to MED12 UF mutations. As noted earlier, TGF-β signalling via SMAD3 directly contributes to alternative splicing, and transcript variants of TGF-β receptors display different binding affinities to TGF-β ligands^48,49^. TGFBR2 (TGF-β type II receptor) for example, has two alternatively spliced variants, TβR-II and TβRII-B; TβR-II only binds to isoforms TGFB1/3 while TβRII-B binds to TGFB2^48,49^. Intriguingly, the dominant transcript usage of TGFBR2 was attributed to TβRII-B (ENST00000359013) in the endometrium from patients with MED12-UF mutant fibroid (Fig. 5d). Notably, most genes identified in the DTU analysis did not exhibit differential expression at the gene level, with only 174 genes overlapping in both analyses (Fig. 5e). Additionally, besides angiogenesis, genes involved in prostaglandin synthesis (PTGES, PTGES2, and PTGFR), progesterone receptor (PGR), and FGF signalling (FGF7 and FGFR2) were identified solely through the DTU analysis (Fig. 5e).

**Fig 5:**
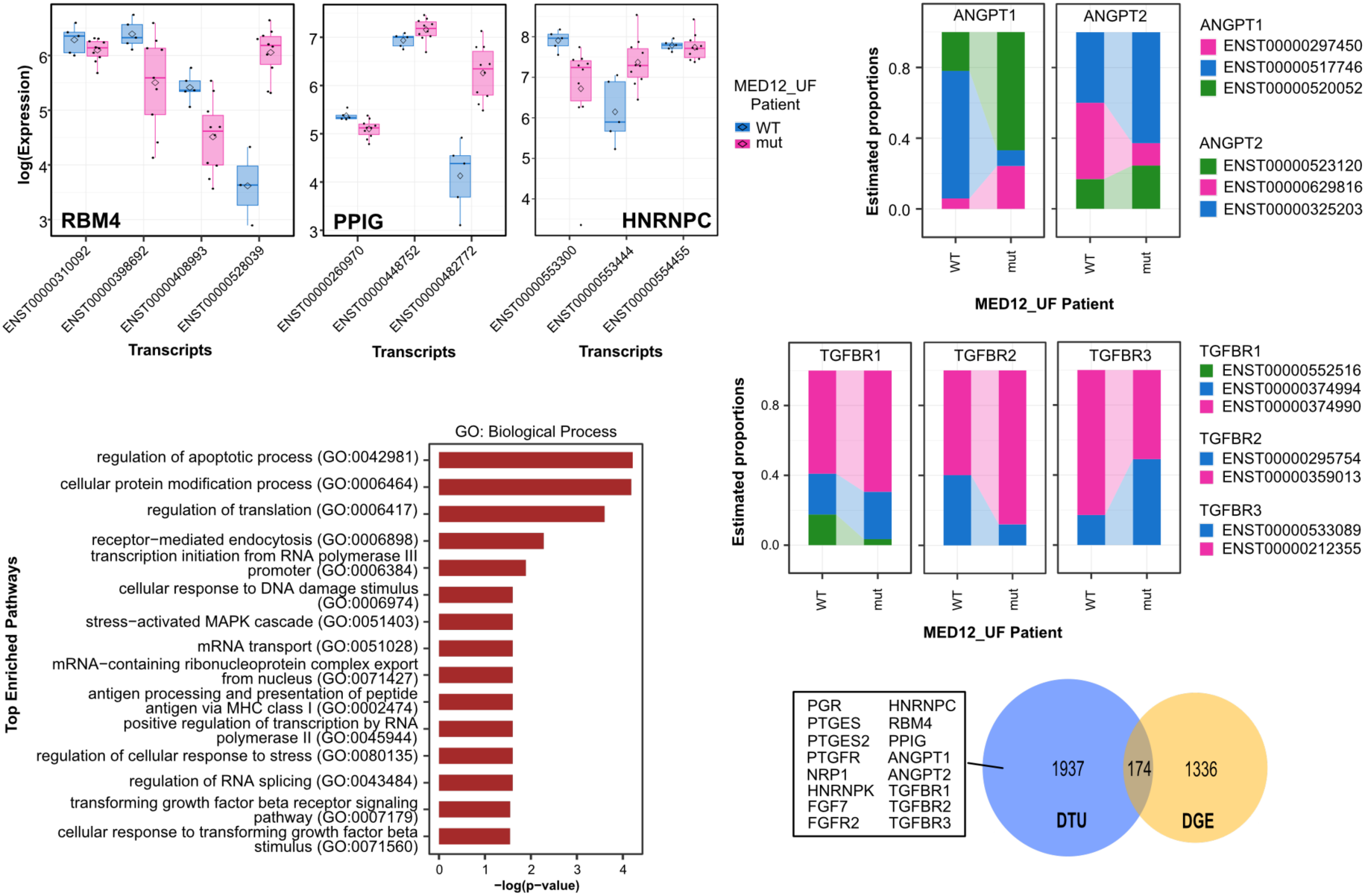
Differential transcript usage analysis of the endometrium samples from patients with MED12-UF mutated or WT fibroids. **a)** Expression of transcript variants in the genes involved in RNA splicing. **b)** The enriched pathways of genes with differential transcript usage detected by DRIMSeq. **c, d)** Ribbonplot demonstrates the changes of transcript usage between the conditions. **e)** Venn diagram reveals differences between differential expressed genes and differential transcript usage in the endometrium samples acquired from patients with MED12-UF mutated or WT fibroid.

### Single-cell transcriptomic analysis identifies altered TGF-β signalling and ECM in UF endometrium and myometrium

The impact of fibroids on endometrial function has been reviewed by Ikhena and Bulun (2018)^50^. Elevated TGF-β3 level secreted by fibroids may hinder expression of genes in wound healing and coagulation pathways, potentially contributing to HMB^51^. To gain insights into differences between UF and healthy endometrium, we conducted single-cell RNAseq on endometrium from UF patients with HMB, integrating and comparing them with healthy endometrium at secretory phase^52^. Following successful batch correction, quality control and cell annotation (cell annotation shown in Wang et al, 2024, under revision), 6 major cell types were further detailed into 10 cell clusters, including lymphatic endothelial cells, macrophage, and dendritic cells. Quantitative differences were observed in several clusters between normal and diseased tissue (Wang et al, 2024). Leveraging CellChat^53^ for ligand-receptor interaction analysis in single-cell datasets (Fig. 6), we observed significantly increased cross-talks between UF-patient derived endometrial clusters, compared to healthy endometrium (Fig. 6a and c, left panel). Multiple signalling pathways including TGF-β and BMP signalling (Fig. 6a), were enriched and the expression of receptors for these pathways (TGFBR1, TGFBR2, BMPR1A, BMPR1B, BMPR2 and ACVR1) markedly increased in UF-patient endometrium (Fig. 6b). Similarly, our integrated analysis (Supplementary Fig. 20) of single-cell myometrium datasets^54,55^ showed elevated signalling pathways in UF-patient myometrium, including uniquely existing pathways such as TGF-β (right panel in Fig. 6c). The receptors of TGF-β were highly expressed in UF-patient myometrium, compared to the normal ones (Fig. 6d). The previously established high levels of TGF-β in uterine fibroid tissue^56,57^ indicates fibroid as the likely source of TGF-β ligands, suggesting a critical role of TGF-β signalling and potential communication between fibroid and uterine tissues. Cross-talk differences in collagen and laminin signalling between normal and UF tissues (Fig. 6 and Supplementary Fig. 21) implicate possible distinctions in cellular microenvironment, ECM and basement membrane architecture in uterine tissues between normal and UF patients. Dysfunctional ECM due to fibroid formation may subsequently impair homeostasis of endometrium and myometrium.

**Fig. 6:**
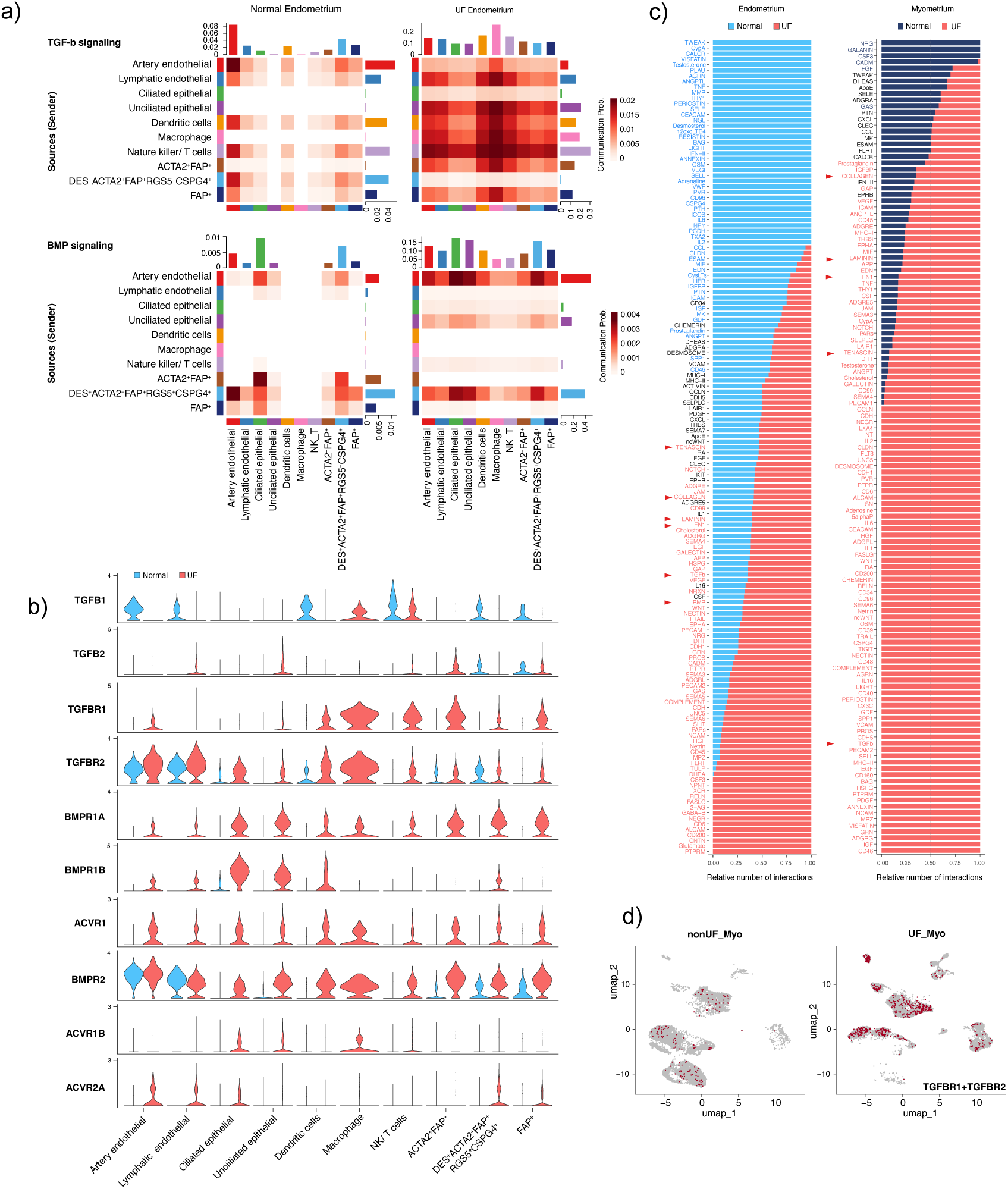
Single-cell transcriptomics of tissues obtained from UF and normal donors. **a)** Heatmap of TGF-β (upper panel) and BMP signaling (bottom) among the clusters of UF and normal endometrium. **b)** Expression levels of ligands and receptors involved in TGF-β and BMP signaling, in normal (blue) and UF (red) endometrium, shown as Vlnplot. **c)** Stacked barplot showing relative ratio of interactions in each signaling pathway predicted in the endometrium (left) and myometrium (right) obtained from normal (blue) and UF (red) donors. Triangles in red highlight the signalling pathways known to be elevated in leiomyoma. **d)** Feature plot showing the expression of TGFBR1 and TGFBR2 in normal (left) and UF (right) myometrium.

### TGF-β signalling in THESC cells induces alternative splicing

As described above in the bulk short-read RNAseq data, TGF-β signalling may induce significant changes in alternative splicing in uterine tissues. To probe potential switches in transcript usage induced by TGF-β signalling within endometrium, we treated the THESC (h<u>T</u>ERT-immortalized <u>H</u>uman <u>E</u>ndometrial <u>S</u>tromal <u>C</u>ells) cell line with TGF-β during *in vitro* decidualization and monitored transcript-level changes using Nanopore long-read RNA sequencing (Fig. 7 and Supplementary Fig. 22) allowing unambiguous determination of transcript isoforms. A MAPK/ERK kinase (MAP2K, MEK) inhibitor (MEKI)^58^ was applied to probe the effect of TGF-β by blocking the downstream signalling cascade. Consistent with our short-read (Illumina sequencing) dataset of THESC decidualization, the enriched pathways of differentially expressed genes (padj < 0.05, abs(log2FC) ≥ 1.5) include cell cycle and chromosome segregation (Supplementary Fig. 23). We employed Talon^59^ and Swan^60^ to identify and quantify transcripts isoforms in the long-read dataset. Importantly, TGF-β treatment during the decidualization led to differential transcript usage, particularly in genes involved in mRNA processing and splicing such as the hnRNP family^61–63^ (HNRNPA1, A2B1, C, K, R, U), RNA-binding proteins (RBM4, RBM39)], VEGFA-VEGFR2 signalling pathways, and a geneset related to hereditary leiomyomatosis (Supplementary Fig. 24a), compared to the treatment using DMSO as solvent control. Similar pathways were also enriched when comparing the treatment of TGF-β plus MEKI or TGF-β versus control conditions (Supplementary Fig. 24b). As shown in Fig. 7, the treatments significantly altered the ratio of transcripts for several genes. For instance, HNRNPA2B1 showed a shift from 100% A2B1-202 to 50% of A2B1-202 and A2B1-206 upon TGF-β treatment. While A2B1-206 is an intron-retained, non-protein coding transcript, this potentially signifies down-regulation of HNRNPA2B1. In addition to A2B1, 10 HNRNPC transcript isoforms were detected (Fig. 7a); aside from HNRNPC-206 (ENST00000553444) as an intron-retained, non-protein coding transcript, the remaining isoforms are all protein-coding, with changes in protein structure which may be associated with altered function. The altered ratio of these transcript isoforms may indicate that expression and function of hnRNP family members are regulated via the transcript switches and subsequently affect RNA splicing and processing.

**Fig. 7:**
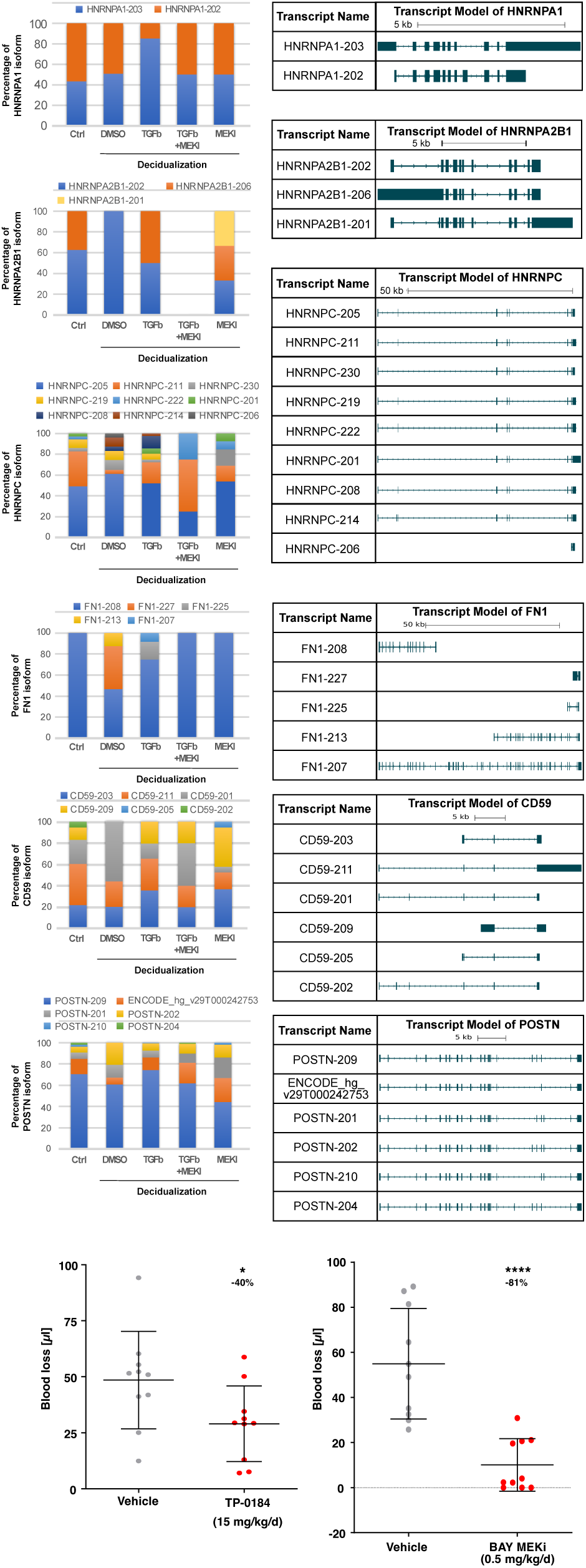
Switches of transcript usage under TGF-β treatment during *in vitro* decidualization, identified by Nanopore long-read sequencing. Transcript isoforms in THESC cell line treated with **(a)** TGF-β or **(b)** TGF-β plus MEK inhibitor during decidualization **a)** Percentage of isoform expression of three splicing factors, HNRNPA1, HNRNPA2B1, and HNRNPC. **b)** Percentage of isoform expression of FN1, CD59 and POSTN. **c)** MEKI (left) or ACVR inhibitor (TP-0184, right) treatment leads to abrogation of menstrual-like bleeding in a murine model. Total uterine blood loss measured by alkaline elution of tampons corrected by background (n = 10 per experiment). Data are shown as mean with SD and significance tested by Students t test (* p<0.05, **** p<0.0001).

In addition to the hnRNP family, transcript switches in genes involved in ECM structure were observed (Fig. 6b). FN1 (fibronectin), a key glycoprotein of the ECM structure, mediates multiple cellular processes through its integrin-, collagen-, and growth factor (i.e. TGF-β)-binding domains^64–66^. Subject to alternative splicing, fibronectin has multiple isoforms that may be composed of the N-terminal, central- and C-terminal domain, in addition to EDA, EDB. The resulting protein products from alternatively spliced transcripts exhibit differential ligand-binding affinities, as well as different properties regarding dimerization, solubility and fibrillogenesis^64,65^. Among the protein coding transcripts we identify FN1-208 (encodes 73 kDa N-terminal protein), FN1-213 (encodes about 121 kDa protein composed of central and C-terminal domains) and FN1-207 (encodes about 239 kDa full-length protein containing V64 variant of the V region, without EDA and EDB regions). These variants may execute different protein functions that subsequently alter the ECM organisation. Another ECM protein, POSTN (periostin), a secreted ECM glycoprotein involved in cell adhesion, has been found to be up-regulated in multiple pathological processes such as fibrosis and tumour progression^67^. For the POSTN transcripts detected in the THESC cells, the sites of alternative splicing are mainly located at the regions of exon 17 to exon 21 (Fig. 6b) and the expression of these isoforms has been reported in normal and diseased tissues^67^.

We correlate the MEKI and TGF-β inhibitor treatments using *in vivo* studies in a mouse menstruation model^68^. This *in vivo* murine system mimicks a primate menstrual cycle and bleeding using ovariectomised mice which are primed with estrogen, progesterone and a physical stimulus within the uterus which is followed by progesterone withdrawal-induced menstrual-like bleeding. We observe reduced uterine bleeding following MEK or ACVR1(blocking TGF-β receptor) inhibition (Fig. 7c). Combined, the data support a hypothesis where UF-associated growth factors and cytokines lead to alternative splicing in the endometrium, which may be associated with HMB.

Overall, our results show transcript switches in the decidualized THESC cell line and the UF endometrium from patients with MED12-UF mutations.

## DISCUSSION

Our study highlights several novel aspects in uterine fibroid biology. First, our targeted sequencing approach towards known genetic drivers of UF, which include mutations in the MED12 and HMGA2 genes, as well as less common mutations in other genes including FH ^14,15^, COL4A5/6 ^11,16^, HMGA1 ^32^, RAD51B ^11^, AHR ^33^, CAPRIN1 ^33^, CUX1 ^34^, DCN ^33^ and PCOLCE ^35^. Unexpectedly our findings reveal a greater proportion than previously reported of missense, insertion-deletion and frameshift variants of COL4A6 that may impact protein functions, in addition to variants of COL4A5 and COL4A6 that are genetically associated with AS-DL. Conversely, we identify a significantly lower proportion of mutations and structural variants in MED12 and HMGA2 (<50% of cases in our study), previously shown to account for ∼90% of disease cases. The reason for these discrepancies is unknown but the data may point to significant ethnic and regional differences in genomic aberrations found in UFs.

The results of our study extend previous findings of extracellular matrix (ECM) proteins in UF. Excessive deposition of ECM is a hallmark of fibrotic diseases and has been linked to abnormal HMB and pelvic pressure and pain ^69,70^. The ECM is a complex network of over 300 proteins, including collagens, proteoglycans, and glycoproteins, as well as many ECM-modifying proteases, growth factors, chemokines, and accessory proteins ^71^ playing a key role in cell proliferation, migration, morphogenesis, tissue homeostasis and communication with the surrounding microenvironment. Studies have demonstrated a direct link between excessive ECM production and UF pathogenesis, with events such as the menstrual cycle and decidualization shown to trigger an inflammatory response that promotes ECM production and repair ^72–75^. Expectedly, our transcriptomic and proteomic data confirm an increase in the expression of COL1A1, COL3A1 and VCAN in fibroid tissue compared to myometrium, in line with previous studies which demonstrate that both the mRNA and mature cross-linked protein collagen are overexpressed in UFs ^21,76–80^. We observed elevated receptor-ligand interactions in collagen and laminin signalling in the single-cell datasets of endometrium and myometrium from UF patients, highlighting the dynamic nature of ECM in uterine tissues and pointing towards a role of basement membrane architecture in patient endometrium associated with fibroids ^31,81^. Our findings, in conjunction with other studies, strongly suggests that UFs are characterised by disrupted ECM composition and mechanical signalling, which are believed to play major roles in their growth and development. As current treatments for UFs are limited, targeting ECM-related pathways may offer avenues for treatment, as evidenced by ongoing studies ^82–85^.

Second, we are providing evidence that gene and protein expression profiles in the endometrium are different when linked to the presence of fibroids; this observation may provide insights into potential mechanisms for associated abnormal bleeding. We identified gene profiles that are dominated by RNA metabolic processes and RNA splicing. Alternative splicing, the process by which exons of a gene are included or excluded in the mRNA as template for protein translation, allows one gene to encode multiple proteins, often with distinct biological function, thereby acting as a major contributor to protein diversity^86^. Alternative splicing is linked to various diseases, including cancer^87^. The spliceosome complex, composed of proteins such as DDX46, PRPF6, SNRNP70, SNRPD2, SRRM2, plays a key role in this process by cutting out introns from pre-RNA. In our analysis of endometrium samples, we detect that these spliceosome components itself are differentially expressed via transcript switches, suggesting a potential role in endometrial physiology, akin to similar findings in diseases like Alzheimer’s and cancer^88,89^. Our multifactorial analysis strategy also identifies a difference in RNA processing and splicing correlated with MED12 mutational status and association with HMB. The regulation of RNA splicing is a complex process that can be regulated by various growth factors and cytokines, such as TGF-β and EGF^90,91^, through signal transduction pathways such as SMAD3, PI3K/Akt/SRPK1, and others^92–97^. These signalling pathways have been shown to directly influence alternative splicing of various genes. At the single-cell level, we see that signalling events in the endometrium and myometrium of UF patients are higher than in healthy individuals; several pathways, including TGF-β and BMP, are differentially enriched in UF patient endometrium and myometrium. The observed high expression of the receptor genes for these signalling molecules in UF patient endometrial and myometrial cells suggests that those patient cells show increased responses to TGF-β and BMP signalling molecules, which likely originate from the fibroid tissue itself. As a proof of concept that splicing plays a significant role in endometrial tissues, we investigated transcript isoform changes in endometrial stroma cell cultures during decidualisation. Indeed, we identify differential transcript expression of members of the hnRNAP family of RNA binding proteins which may be associated with changes in transcript isoforms we also observe for complement receptor CD59, besides components of ECM such as periostin and fibronectin that contribute to endometrium physiology^98–103^. These alternative transcript isoforms may indeed play a significant role in regulating properties of different cell adhesion and ligand-binding affinities and accordingly may impact ECM organisation in the homeostasis of endometrium. Notably, ECM stiffness has previously been linked to alternative splicing events by activation of Ser/Arg-rich protein components of the spliceosome^104^. Lastly, we show that effective in vivo approaches such as inhibition of MEK and TGF-β signalling in a pre-clinical model to reduce HMB can be linked in the THESC model with alternative splicing events. Taken together, the integrated systems approach has uncovered novel genomic facets associated with uterine fibroids, provided integration of transcriptomic and proteomic data highlighting the ECM as critical factor in leiomyoma pathobiology and highlights alternative splicing as a significant contributor to fibroid biology.

## AUTHOR CONTRIBUTIONS

UO, CYW, MP and APC designed and supervised the study; CYW, MP and UO wrote the first manuscript draft. KZ, CMB, JM, KG, SM, MM supervised and performed sample collection and clinical annotation, with important help from CMB, TMZ and ALH. CYW, MP, DOB, JM, NM, VG, BM, SB, RF performed experiments. CYW, DOB, JM, APC performed data analysis, with significant contributions from AN, MO, BK, and ALH. CMB, KZ, ALH, SM, JM, NS and TMZ contributed critical data interpretation. All authors have read and provided input to the manuscript.

## ACKNOWLEDGMENTS

This work was supported through the Bayer - Oxford Alliance in Women’s Healthcare, which receives funding through the NIHR Biomedical Research Centre, the Endometriosis CaRe Centre Oxford, Oxford University Medical Sciences Division and Bayer Healthcare. Further research support was obtained from Innovate UK (UO, MP, APC), the National Institute for Health Research Oxford Biomedical Research Centre (UO), Cancer Research UK (CRUK, UO), the Bone Cancer Research Trust (APC and UO), the Leducq Epigenetics of Atherosclerosis Network (LEAN) program grant from the Leducq Foundation (UO), the Chan Zuckerberg Initiative (APC) and the Myeloma Single Cell Consortium (UO). APC is a recipient of an MRC Career Development Fellowship (MR/V010182/1). Work in the BMK laboratory was supported by the Wellcome Trust (097812/Z/11/Z) and the Engineering and Physical Science Research Council (EP/N034295/1).

## Methods

### Patient samples and tissue collection

Fibroid, myometrium, and endometrium tissues were collected from 137 donors undergoing hysterectomy, myomectomy or TransCervical Resection of Fibroids (TCRF) at the John Radcliffe Hospital, Oxford, in accordance with ENDOX study guidelines (09/H0604/58). All experimental protocols were approved by the local Research Ethics Committee (National Health Services (NHS) Research (NRES) Committee South Central-Oxford). Informed written consent was provided by patients participating in the study. In all cases, UF diagnosis was confirmed surgically and by histology. HMB status and use of hormone therapy was established from clinical notes and donor questionnaires. Menstrual cycle phase was determined by histopathology of the endometrium. Tissue samples were collected immediately after surgery, snap frozen in liquid nitrogen, and stored at -80°C.

### SureSelect targeting sequencing

DNA for SureSelect assays and SNP arrays was purified from fresh frozen samples stored at -80°C using a PureLink Genomic DNA Kit (Invitrogen) according to the manufacturer’s instructions for mammalian tissue. Eluted DNA was quantified by NanoPhotometer (Implen) and stored at -20 °C until further use. Approximately 100 ng of each DNA sample was used to create Illumina sequencing libraries using a NEBNext Ultra II FS DNA Library Prep Kit (New England Biolabs (NEB), E7805S). After PCR amplification with index primers, targeted DNAs were captured and enriched by SureSelect XT HS Target Enrichment Kit ILM Hyb Module according to the manufacturer’s instructions (Agilent). Indexed libraries were quantitated by high sensitivity DNA ScreenTape assay for TapeStation (Agilent), pooled at equimolar concentration, and sequenced on a NextSeq 500 to an average of ∼8 million reads/sample. Reads were initially assessed for quality using FastQ Screen v0.14.0, FastQC v0.11.9 and MultiQC v1.5.dev0. Raw reads of each sample were mapped to hg38 using BWA v0.7.17 and merged into a single bam file. For SNPs and small insertions/deletions (indels), variant calling was done by mpileup provided by bcftools v1.9^105^ using human genome GRCh38 to detect variants; variant annotation, effect prediction and associated phenotypes were performed by SnpEff ^106^ and Ensembl Variant Effect Predictor^39^. Structure variants with large deletions or translocations were identified by Gridss v2.13.2^107^.

### Genotyping of SNP arrays

For each genotype batch, samples with a call rate of less than 98% were excluded. Additionally, SNPs with call rate less than 95% were excluded. The batches were then merged into one genotype dataset. The merged genotype data was then imputed up to the Haplotype Reference Consortium (HRC r1.1 2016). In total 8,642,416 SNPs with an imputation R2 > 0.8 and minor allele frequency (MAF) > 0.01 were carried forward for cis-eQTL mapping.

### Whole genome sequencing by Nanopore

DNA for Nanopore sequencing was purified from frozen samples stored at -80 °C using a Blood & Cell Culture DNA Midi Kit (Qiagen). Approximately 100 mg of tissue sample was crushed in a BioPulveriser cooled in liquid nitrogen, before following the manufacturer’s protocol for tissue samples. Eluted DNA was quantified by NanoPhotometer (Implen) and the size and quality assessed by Genomic DNA ScreenTape Analysis for TapeStation (Agilent). Samples were stored at -20 °C until further use. DIN values were >9 and size was >60,000 bp. Whole genome sequencing libraries were generated using ONT Ligation Sequencing Kit V14 (SQK-LSK114) and sequenced on the Nanopore PromethION platform.

### Bulk RNA-sequencing

Tissue samples stored at -80 °C were cyromilled with Trizol without allowing the tissue to thaw. Briefly, one stainless steel end cap was inserted into a polycarbonate cylinder and precooled in liquid nitrogen along with the other cap and impactor. On dry ice, the impactor, 1.6 mL of Trizol and the tissue sample were added to the cylinder, which was capped and placed in the cyromill. The procedure was performed for 3 cycles of 2 min. Once completed, samples were transferred to a 50 mL centrifuge tube pre-chilled on dry ice. When processing multiple samples, tubes were kept on dry ice or stored at -80°C prior to downstream batch processing. Sample tubes were placed in a 37°C water bath until thawed, vortex mixed, aliquoted into 1.5 mL centrifuge tubes and stored at -80°C if not proceeding immediately to RNA extraction. RNA extraction was performed using a Direct-zol RNA miniprep kit (Zymo Research) and on-column DNAse I digest, according to the manufacturer’s instructions. Eluted RNA was quantified by NanoPhotometer (Implen), quality checked by high sensitivity RNA ScreenTape assay for TapeStation (Agilent), and stored at -80°C until further use. RIN values generally ranged between 3 to 5, typical of tissue samples, but suggesting some 3’ bias would be observed in the RNAseq.

Approximately 100 ng of each RNA sample was used to create Illumina sequencing libraries using a NEBNext Ultra II Directional RNA Library Prep Kit for Illumina with NEBNext Poly(A) mRNA Magnetic Isolation Module (New England Biolabs) according to the manufacturer’s instructions. Indexed libraries were quantitated by high sensitivity DNA ScreenTape assay for TapeStation (Agilent), pooled at equimolar concentration, and sequenced on a NextSeq 500 to an average of ∼20 million reads/sample.

### Analysis of bulk RNA-sequencing

Reads were initially assessed for quality using FastQ Screen v0.14.0, FastQC v0.11.9 and MulitQC v1.5.dev0. Raw reads of each sample were then merged into a single file and pseudo-aligned to the human genome hg38 with Kallisto 0.46.0. The samples with alignment rate lower than 60% were excluded from downstream analysis. Using the count matrix produced by Kallisto, differential expression analysis was performed by DESeq2 v1.35.0^108^ for comparisons with the clinical factors such as cycle phase, HMB, MED12 status, and the technique factor like batch effect. Functional analysis including gene set enrichment analysis (GSEA) and over-representation analysis (ORA) were done by R packages clusterProfiler 4.2.2 ^109,110^. For differential transcript usage analysis, raw reads of the samples were pseudo-aligned to gencode.v29.annotation.gtf by Kallisto, and the output abundance files were imported by tximport^111^ and then analysed by DRIMSeq^112^ and DEXSeq^113^.

### Analysis of gene fusion events using bulk RNA-seq dataset

The fastq files for each fibroid sample from bulk RNA sequencing were merged into a single file and aligned to the human genome hg38 with STAR v2.7.6a together with the following settings: --outReadsUnmapped None --outSAMtype BAM Unsorted -- outFilterMultimapNmax 1 --outFilterMismatchNmax 3 --chimSegmentMin 10 --chimOutType WithinBAM SoftClip --chimJunctionOverhangMin 10 --chimScoreMin 1 -- chimScoreDropMax 30 --chimScoreJunctionNonGTAG 0 --chimScoreSeparation 1 -- alignIntronMax 100000 --alignSJstitchMismatchNmax 5 -1 5 5 –chimSegmentReadGapMax 3. The main output files (aligned.out.bam) were then used as inputs in Arriba 1.0.0^41^.

### Uterine fibroid protein extraction

Frozen UF samples were cryomilled on liquid nitrogen in 1.6 mL of a lysis buffer comprising 6M urea, 2M Thiourea, 50% RIPA, 4% SDS, 100 mM DTT, and supplemented with protease and phosphatase inhibitors. To release protein bound to RNA and DNA, 1 μL of benzonase nuclease was added to 500 µL of each thawed sample and incubated on ice for 20 min. Due to the inherent toughness of the UF tissue samples, each was subjected to three rounds of bead beating for 2 min at 4°C for maximum tissue disruption. Samples were then spun down for 5 min at 10,000 g and 4°C. The supernatant was diluted (1:5) in water to achieve a final DTT concentration of 20 mM. Reduced samples were alkylated by adding IAA to a final concentration of 40 mM and incubated at room temperature for 1 hour in the dark. To remove SDS and other contaminants, all samples were subjected to a protein extraction procedure of alternating washes in methanol, chloroform and water. To maximise protein recovery, precipitated pellets were resuspended in 500 µL of 100 mM TEAB buffer, sonicated on ice for 5 min in a water bath, and vortexed at room temperature for 30 min. The protein content of each UF sample was then determined using a standard BCA assay.

### Sample digestion, clean-up, and TMT-labelling

Samples were digested in a 96-well format using the SMART Digest kit provided by Thermo Fisher Scientific. Briefly, 150 µg of each lyophilized UF sample was resuspended in 50 µL of 100 mM TEAB and added to 150 µL of the accompanying SMART Digest buffer. Frozen SMART Digest PCR strips containing immobilized trypsin beads were thawed and spun down at 1000 rpm for 1 min, and at 4°C. Samples (200 µL) were transferred into the appropriate PCR tube and incubated on a heated shaker for 180 min at 70°C and 1400 rpm. Upon completion, samples were spun down at 1000 rpm for 1 min. UF digests were cleaned-up with the aid of a vacuum manifold using the SOLAμ Solid-Phase Extraction (SPE) Plates provided with the kit. Samples were loaded in a 1:1 ratio (v:v) with 0.1% TFA, followed by one wash with 0.1% TFA. Peptides were eluted with 70% ACN into a 96-well collection plate and lyophilised to completion. For TMT-labelling, samples (∼150 µg) were resuspended in 100 µL of 100 mM TEAB. Approximately 10% of each sample was removed for the preparation of global pooled samples. For this, two concentrations were prepared to be included in each TMT 10plex labelling reaction, one undiluted pool of all samples (1X Pool), and a five times diluted pool samples (5X Pool). Immediately before use, TMT label reagents were equilibrated to room temperature. To each 0.8 mg vial, 82 μL of anhydrous acetonitrile was added and the reagent allowed to dissolve for 5 min with occasional vortexing, before being gently centrifuged to gather the solution. For each TMT labelling reaction, 41 μL of the TMT label reagent was added to each 100 μL of UF sample. The reaction was allowed to proceed for 1 hour at room temperature before being quenched for 15 min with 8 μL of a 5% hydroxylamine solution. For each TMT 10plex experiment, an equivalent volume (140 μL) of sample was combined, resulting in a total protein amount of approximately 1.5 mg in a final volume of 1.4 mL. Each concatenated sample was desalted on a C18 solid-phase extraction cartridge (Sep-Pak Plus, Waters).

### High-pH reversed-phase pre-fractionation

Approximately 1.5 mg of digested TMT-labelled material was subjected to off-line high-pH reversed-phase pre-fractionation using the loading pump of a Dionex Ultimate 3000 HPLC with an automated fraction collector and a XBridge BEH C18 XP column (3 × 150 mm, 2.5 μm pore size, Waters no. 186006710). Peptides were separated over a 100 min gradient using two basic pH reversed-phase buffers (A: ammonium hydroxide in 100% water, pH 10; B: ammonium hydroxide in 90% acetonitrile, pH 10). The gradient consisted of a 12 min wash with 1% B, then increasing to 35% B over 60 min, with a further increase to 95% B over 8 min, followed by a 10 min wash at 95% B and a 10 min re-equilibration at 1% B. The flow rate was set to 200 μL/min, with fractions collected every 2 min throughout the run. In total, 50 fractions were collected over the run, but samples were concatenated down to a final of 10 fractions by combining every 10th sample. Each fraction was dried down and resuspended in 30 μL of 2% ACN:0.1% formic acid for analysis by LC–MS/MS.

### High Performance Liquid Chromatography Tandem Mass Spectrometry (LC-MS/MS)

LC-MS/MS analysis was performed using a Dionex Ultimate 3000 nano-ultra high pressure reversed-phase chromatography system coupled on-line to a Q Exactive High Field (HF) mass spectrometer (Thermo Scientific). Samples were separated on an EASY-Spray PepMap RSLC C18 column (500 mm x 75 µm, 2 µm particle size; Thermo Scientific) over a 60 min gradient of 2-35 % acetonitrile in 5 % DMSO, 0.1% formic acid and at 250 nL/min. The mass spectrometer was operated in data-dependent mode for automated switching between MS and MS/MS acquisition. Full MS survey scans were acquired from *m/z* 400-2,000 at a resolution of 60,000 at *m/z* 200 and the top 12 most abundant precursor ions were selected for HCD fragmentation. The resolution of MS2 fragment ion detection was also set to 60,000. Fractions were loaded with adjusted sample volumes to analyze ∼1 μg on column.

### Proteomics Data Analysis

MS raw data were searched against the UniProtKB human sequence data base (92,954 entries) and TMT 10plex quantitation performed using Proteome Discoverer software (v 2.3; Thermo Scientific). Search parameters were set to include carbamidomethyl (C) as a fixed modification, with TMT 6plex, oxidation (M), and deamidation (NQ) set as variable modifications. A maximum of 2 missed cleavages were allowed. TMT 10plex quantitation and data analysis was performed in Perseus (v1.6.0.2), resulting in the generation of hierarchical clustering, principal component analysis, and Volcano plots. For PCA analysis, samples underwent log2 transformation and all missing values were removed. This was then followed by median subtraction normalization. For generation of volcano plots, an identical processing workflow was used, but only 50% of missing values were removed. The missing values that remained were imputed from the normal distribution (width 0.3, down shift 1.8). Differentially regulated proteins between groups of interest were subject to gene ontology and pathway enrichment analysis using STRINGdb (https://string-db.org/). Shortlisted targets were further assessed for their biological relevance and therapeutic potential in the treatment of UFs using TargetDB (https://pypi.org/project/targetDB/).

### Integration of transcriptomics and proteomics by Multi-Omics Factor Analysis (MOFA)

The log-normalized count matrices of transcriptomics and proteomics were used as the input data to MOFA ^43,44^, in addition of the metadata containing the clinical information related to donors. MOFA object was prepared by using the default options and trained with the number of factors suggested by MOFA. Donor parameters used in the analysis for factor correlation include genotypes identified by SureSelect targeting sequencing such as MED12 status (WT or mutant at canonical UF mutation) and SNPs associated with COL4A5 GWAS phenotypes, fibroid occurrence (with or without UF), tissue type (such as UF or myometrium), cycle phase (proliferative, secretory, mense, or inactive), and HMB symptom. GSEA analysis was performed by the function implemented in MOFA, using all features associated with factors as input. Selected features with weight value higher than a cut-off value of 0.3 were selected for pathway analysis via an R interface to the Enrichr database^114^ and visualized by STRINGdb.

### Nuclei preparation for single cell RNAseq

A petri dish, 50 ml centrifuge tube, scalpel and forceps were precooled on dry ice before pseudocapsule samples were removed from -80°C and placed in the petri dish. Typical sample sizes ranged from 100-500 mg. Tissue was cut into thin slices and transferred to centrifuge tubes. If processing multiple samples, cut tissue could be stored at -80°C until use. Sample tubes were transferred to wet ice, 4 ml of ice-cold CST buffer (146 mM NaCl, 10 mM Tris-HCl pH 7.5, 1 mM CaCl, 21 mM MgCl2, 0.5% CHAPS (w/v), 0.01% BSA (w/v), 4 ul/ml SUPERaseIN, 4 ul/ml RNasein Plus, 1 cOmplete protease inhibitor tablet/10 ml) added and tubes placed on a rotator for 10 minutes at 4°C. Samples were passed through 30 µm cell strainers (MACS SmartStrainer) into prechilled 15 ml collection tubes on ice. Sample tubes were rinsed with 2 ml ice cold PBS + 1% BSA, which was added to the cell strainer. Cell strainers were rinsed with an additional 2 ml ice cold PBS + 1% BSA and collection tubes centrifuged at 500g for 5 minutes at 4°C. Supernatant was removed and pellet washed by resuspending in 10 ml ice cold PBS + 1% BSA, centrifugation at 500g for 5 minutes at 4°C, removal of supernatant and resuspension in 500 µl ice cold PBS + 1% BSA. A subsample of the nuclei preparation was incubated with DAPI (1 µg/ml) for 5 minutes, added to a haemocytometer and counted under a fluorescent microscope. Concentration of the nuclei was adjusted to ∼1,000 cells/µl and used as input for analysis by 10X Chromium single cell gene expression.

### Library preparation and Sequencing of single-cell RNA sequencing

Chromium single cell gene expression (10X Genomics) was performed using the Chromium Next GEM Single Cell 3’ GEM, Library & Gel Bead Kit v3.1, Chromium Next GEM Chip G Single Cell Kit and Single Index Kit T Set A according to the manufacturer’s instructions starting with 20,000 nuclei as input. Resulting libraries were quantitated by TapeStation (Agilent), pooled at equimolar concentration and sequenced (Novogene (UK) Ltd or Genewiz GmbH) on an Illumina NovaSeq 6000 using a S4 Reagent Kit v1.5 to give ∼30,000 reads/cell.

### Analysis of Single cell RNA sequencing

Raw sequencing data (fastq files) were processed using the scflow workflows (https://github.com/Acribbs/scflow). The Kallisto BUS/BUStools (v0.39.3) workflow1 was implemented to pseudo-align the reads, with a K-mer size of 31 base pairs. Homo sapiens (human) genome assembly GRCh38 (hg38) was used to construct a reference transcriptome. Individual samples of single-nuclei or single cells were analyzed by the pipeline of quantnuclei or quantcells implemented in the scflow workflows, respectively. The output was converted to single-cell experiment objects^115^ and then to Seurat objects (Seurat v4.0)^116^. Quality control and filtering was performed on the Seurat objects; any cell with a mitochondrial ratio higher than 0.1, or fewer than 300 features were removed. Doublets in the samples were detected using the R package scDblFinder^117^ and removed in the scflow pipeline with Seurat clustering.

To integrate the endometrium samples with the published data, we first used the VST method provided by Seurat for variable gene selection and applied Harmony v1.04^118^ for batch correction. Highly variable genes that account for cellular heterogeneity in each main cluster were used and cells were aligned using Harmony. For cell-cell communication, we applied CellChat (v1.4.0)^53^ with input of two matrices, log normalized count matrix and a matrix of the cell label.

### THESC decidualization

THESC cell line was incubated in DMEDM/F-12 with bicarbonate and HEPES (Sigma Cat# D 2906) supplemented with 10% fetal bovine serum (FBS, Charcoal stripped F6765-500ML), puromycin (500 ng/ml), and 1% ITS Premix Universal Culture Supplement (Corning 354350). For the three days experiment of decidualization, cells were seeded at 6-well plates for 40,000 cells per well and incubated overnight. At the next day (Day 0), the decidualization were induced by adding the following reagents into cell medium: Medroxyprogesterone 17-acetate (Sigma, M1629; final conc. 1.0 µM), E2 (estradiol, final conc. 10 nM; Sigma E1024), 8-Br cAMP (8-Bromoadenosine 3′,5′-cyclic monophosphate, final conc. 500 µM; Sigma B6386-100mg). In addition to stimulation for decidualization, cells were further treated with DMSO as mock, TGF-β (10 ng/ml; Millipore GF346) together with or without MEK inhibitor (BAY 1076672, 100 ng/ml) since Day 0, depending on the experimental design. Cells were harvested on Day 3 using Direct-zol RNA MiniPrep kit (Cambridge Bioscience, R2052).

### Library construction and sequencing of Nanopore long read sequencing

50 ng RNA of each sample were reverse transcribed and barcoded by using the PCR-cDNA barcoding kit (SQK-PCB111.24) and NEBNext Companion Module (NEB E7180L). Libraries were then sequenced on the Nanopore PromethION platform.

### Analysis of long read sequencing to identify transcript isoforms

Base calling of fast5 files was done by Guppy (https://github.com/asadprodhan/GPU-accelerated-guppy-basecalling) and converted to fastq format. Reads of each sample were then aligned to hg38 genome by Minimap2^119^ with - - MD flag enabled and output as SAM format. The SAM file of each sample was processed by TALON^59^ for isoform-level analysis, and visualized by Swan^60^ for transcript switching genes and novel transcripts due to alternative splicing.

### Mice

Female C57Bl/6N mice were obtained from Charles River WIGA GmbH (Sulzfeld, Germany) and were used at 8–10 weeks of age. All animal experiments were performed under standardized conditions and in accordance with institutional, state and federal guidelines. The study was approved by the Regional Office for Health and Social Affairs in Berlin (LAGeSo; protocol A0384/ 09).

### Mouse Model of Menstruation and Treatment Regimens

The mouse model of menstruation was performed as described previously ^68^. One week after ovariectomy female C57Bl/6 mice received s.c. injections of 100 ng 17a-estradiol (E2, internal source) in ethanol/ arachis oil (1:9) on three consecutive days. After a three days break a progesterone (P4) releasing silastic tube (0.5 mg P4/ d, internal source ^120^) was implanted s.c. into the back of mice followed by further applications of 5 ng E2 on three consecutive days. Concomitant with the last E2 treatment 50 ml sesame oil were injected into one uterus horn to induce decidualization. 4 days later the P4 implant was removed to initiate P4 withdrawal. Mice were sacrificed at indicated points of time and uteri were weighed and harvested for further analyses. All surgeries were performed under isofluran-induced anesthesia.

Inhibition of uterine bleeding *in vivo* was performed by p.o. application of MEK inhibitor (BAY MEKi, cpd 26^58^, Bayer AG, Germany) at doses of 0.5 mg/kg daily or ACVR1 inhibitor (TP-0184, Toledo Pharmaceuticals, USA) at doses of 15 mg/kg daily dissolved in N-methyl-2-pyrrolidone (NMP)/ polyethylene glycol 400 (PEG400) (1/9) (d0-d15). Controls were treated with vehicle alone.

### Assessment of Uterine Blood Loss and Bleeding Intensity

To assess bleeding intensity, cotton swabs soaked with 30 ml PBS solution were used to catch up vaginal fluid containing uterine blood. Swabs were pressed onto a glass slide. Dried smears were incubated in 70% ethanol for 2 hours and hematoxylin/eosin (H&E) stained using an automatic Leica Multistainer and Coverslipper (Leica ST5020+CV5030, Leica Microsystems, Wetzlar, Germany). Using a scoring system bleeding intensity was evaluated in vaginal smears by microscopy : 0= no erythrocytes; 1 = few erythrocytes; 2 = evenly scattered erythrocytes; 3= frequent occurrence of erythrocytes with little cluster formation; 4 = very frequent occurrence of erythrocytes, strong cluster formation; 5= massive accumulations and clustering of erythrocytes.

To quantify the total amount of blood loss, tampon-like cotton pads (4–4.8 mm diameter, Roeko, Coltène/Whaledent, Altstä tten, Switzerland) were inserted into the vagina of mice at P4 withdrawal. Mice additionally received a paper collar to prevent removal of tampons. Tampons were changed twice daily and collected for each mouse separately. Blood volume was quantified by the alkaline hematine method reported elsewhere^121^. Briefly, tampons were left to dry at room temperature. Heme chromogens were dissolved in 1000 ml 5% NaOH (w/v) and rotated over night at room temperature. Optical density of the eluate was measured in an ELISA Reader at a wavelength of 546 nm. Blood volume contained in cotton swabs was calculated based on a regression curve of standards prepared from venous blood.

## Data availability

All raw and processed sequencing data are available in the NCBI’s Gene Expression Omnibus: bulk RNA Sequencing data (GSE199849) and single-cell RNA sequencing data (GSE220650) of patient samples applied to this study.

## REFERENCES

1. Baird, D.D., Dunson, D.B., Hill, M.C., Cousins, D. & Schectman, J.M. High cumulative incidence of uterine leiomyoma in black and white women: ultrasound evidence. Am J Obstet Gynecol 188, 100–107 (2003).

2. Gupta, S., Jose, J. & Manyonda, I. Clinical presentation of fibroids. Best Pract Res Clin Obstet Gynaecol 22, 615–626 (2008).

3. Stewart, E.A. Clinical practice. Uterine fibroids. N Engl J Med 372, 1646–1655 (2015).

4. Jacobson, G.F., Shaber, R.E., Armstrong, M.A. & Hung, Y.Y. Hysterectomy rates for benign indications. Obstet Gynecol 107, 1278–1283 (2006).

5. Cardozo, E.R., et al. The estimated annual cost of uterine leiomyomata in the United States. Am J Obstet Gynecol 206, 211 e211–219 (2012).

6. Wegienka, G., et al. Self-reported heavy bleeding associated with uterine leiomyomata. Obstet Gynecol 101, 431–437 (2003).

7. Cooper, K.G., Jack, S.A., Parkin, D.E. & Grant, A.M. Five-year follow up of women randomised to medical management or transcervical resection of the endometrium for heavy menstrual loss: clinical and quality of life outcomes. BJOG 108, 1222–1228 (2001).

8. Goodman, A. Abnormal genital tract bleeding. Clin Cornerstone 3, 25–35 (2000).

9. Hapangama, D.K. & Bulmer, J.N. Pathophysiology of heavy menstrual bleeding. Womens Health (Lond) 12, 3–13 (2016).

10. Makinen, N., et al. MED12, the mediator complex subunit 12 gene, is mutated at high frequency in uterine leiomyomas. Science 334, 252–255 (2011).

11. Mehine, M., et al. Characterization of uterine leiomyomas by whole-genome sequencing. N Engl J Med 369, 43–53 (2013).

12. Elmlund, H., et al. The cyclin-dependent kinase 8 module sterically blocks Mediator interactions with RNA polymerase II. Proc Natl Acad Sci U S A 103, 15788–15793 (2006).

13. Cleynen, I., et al. HMGA2 regulates transcription of the Imp2 gene via an intronic regulatory element in cooperation with nuclear factor-kappaB. Mol Cancer Res 5, 363–372 (2007).

14. Ono, M., et al. Paracrine activation of WNT/beta-catenin pathway in uterine leiomyoma stem cells promotes tumor growth. Proc Natl Acad Sci U S A 110, 17053–17058 (2013).

15. Stewart, E.A., et al. Uterine fibroids. Nat Rev Dis Primers 2, 16043 (2016).

16. Mehine, M., et al. Integrated data analysis reveals uterine leiomyoma subtypes with distinct driver pathways and biomarkers. Proc Natl Acad Sci U S A 113, 1315–1320 (2016).

17. Arslan, A.A., et al. Gene expression studies provide clues to the pathogenesis of uterine leiomyoma: new evidence and a systematic review. Hum Reprod 20, 852–863 (2005).

18. Vanharanta, S., et al. Distinct expression profile in fumarate-hydratase-deficient uterine fibroids. Hum Mol Genet 15, 97–103 (2006).

19. Christacos, N.C., Quade, B.J., Dal Cin, P. & Morton, C.C. Uterine leiomyomata with deletions of Ip represent a distinct cytogenetic subgroup associated with unusual histologic features. Genes Chromosomes Cancer 45, 304–312 (2006).

20. Vanharanta, S., et al. 7q deletion mapping and expression profiling in uterine fibroids. Oncogene 24, 6545–6554 (2005).

21. Leppert, P.C., Catherino, W.H. & Segars, J.H. A new hypothesis about the origin of uterine fibroids based on gene expression profiling with microarrays. Am J Obstet Gynecol 195, 415–420 (2006).

22. Zavadil, J., et al. Profiling and functional analyses of microRNAs and their target gene products in human uterine leiomyomas. PLoS One 5, e12362 (2010).

23. Hodge, J.C., et al. Expression profiling of uterine leiomyomata cytogenetic subgroups reveals distinct signatures in matched myometrium: transcriptional profilingof the t(12;14) and evidence in support of predisposing genetic heterogeneity. Hum Mol Genet 21, 2312–2329 (2012).

24. Cirilo, P.D., et al. An integrative genomic and transcriptomic analysis reveals potential targets associated with cell proliferation in uterine leiomyomas. PLoS One 8, e57901 (2013).

25. Ura, B., et al. Identification of proteins with different abundance associated with cell migration and proliferation in leiomyoma interstitial fluid by proteomics. Oncol Lett 13, 3912–3920 (2017).

26. Ura, B., et al. Two-dimensional gel electrophoresis analysis of the leiomyoma interstitial fluid reveals altered protein expression with a possible involvement in pathogenesis. Oncol Rep 33, 2219–2226 (2015).

27. Liu, Y., Lu, D., Sheng, J., Luo, L. & Zhang, W. Identification of TRADD as a potential biomarker in human uterine leiomyoma through iTRAQ based proteomic profiling. Mol Cell Probes 36, 15–20 (2017).

28. Ko, Y.A., et al. Extracellular matrix (ECM) activates beta-catenin signaling in uterine fibroids. Reproduction 155, 61–71 (2018).

29. Jamaluddin, M.F.B., et al. Proteomic Profiling of Human Uterine Fibroids Reveals Upregulation of the Extracellular Matrix Protein Periostin. Endocrinology 159, 1106–1118 (2018).

30. Jamaluddin, M.F.B., Nagendra, P.B., Nahar, P., Oldmeadow, C. & Tanwar, P.S. Proteomic Analysis Identifies Tenascin-C Expression Is Upregulated in Uterine Fibroids. Reprod Sci 26, 476–486 (2019).

31. Jamaluddin, M.F.B., Nahar, P. & Tanwar, P.S. Proteomic Characterization of the Extracellular Matrix of Human Uterine Fibroids. Endocrinology 159, 2656–2669 (2018).

32. Williams, A.J., Powell, W.L., Collins, T. & Morton, C.C. HMGI(Y) expression in human uterine leiomyomata. Involvement of another high-mobility group architectural factor in a benign neoplasm. Am J Pathol 150, 911–918 (1997).

33. Yatsenko, S.A., et al. Highly heterogeneous genomic landscape of uterine leiomyomas by whole exome sequencing and genome-wide arrays. Fertil Steril 107, 457–466 e459 (2017).

34. Schoenmakers, E.F., et al. Identification of CUX1 as the recurrent chromosomal band 7q22 target gene in human uterine leiomyoma. Genes Chromosomes Cancer 52, 11–23 (2013).

35. Medikare, V., Kandukuri, L.R., Ananthapur, V., Deenadayal, M. & Nallari, P. The genetic bases of uterine fibroids; a review. J Reprod Infertil 12, 181–191 (2011).

36. Landrum, M.J., et al. ClinVar: public archive of interpretations of clinically relevant variants. Nucleic Acids Res 44, D862–868 (2016).

37. Sudmant, P.H., et al. Global diversity, population stratification, and selection of human copy-number variation. Science 349, aab3761 (2015).

38. Sollis, E., et al. The NHGRI-EBI GWAS Catalog: knowledgebase and deposition resource. Nucleic Acids Res 51, D977–D985 (2023).

39. McLaren, W., et al. The Ensembl Variant Effect Predictor. Genome Biol 17, 122 (2016).

40. Miner, J.H. Alport syndrome with diffuse leiomyomatosis. When and when not? Am J Pathol 154, 1633–1635 (1999).

41. Uhrig, S., et al. Accurate and efficient detection of gene fusions from RNA sequencing data. Genome Res 31, 448–460 (2021).

42. Maybin, J.A. & Critchley, H.O. Menstrual physiology: implications for endometrial pathology and beyond. Hum Reprod Update 21, 748–761 (2015).

43. Argelaguet, R., et al. Multi-Omics Factor Analysis-a framework for unsupervised integration of multi-omics data sets. Mol Syst Biol 14, e8124 (2018).

44. Argelaguet, R., et al. MOFA+: a statistical framework for comprehensive integration of multi-modal single-cell data. Genome Biol 21, 111 (2020).

45. Cox, J. & Mann, M. 1D and 2D annotation enrichment: a statistical method integrating quantitative proteomics with complementary high-throughput data. BMC Bioinformatics 13 Suppl 16, S12 (2012).

46. Brodsky, R.A. Paroxysmal nocturnal hemoglobinuria. Blood 124, 2804–2811 (2014).

47. Dickson, K.A., et al. Ribonuclease inhibitor regulates neovascularization by human angiogenin. Biochemistry 48, 3804–3806 (2009).

48. Rotzer, D., et al. Type III TGF-beta receptor-independent signalling of TGF-beta2 via TbetaRII-B, an alternatively spliced TGF-beta type II receptor. EMBO J 20, 480–490 (2001).

49. del Re, E., Babitt, J.L., Pirani, A., Schneyer, A.L. & Lin, H.Y. In the absence of type III receptor, the transforming growth factor (TGF)-beta type II-B receptor requires the type I receptor to bind TGF-beta2. J Biol Chem 279, 22765–22772 (2004).

50. Ikhena, D.E. & Bulun, S.E. Literature Review on the Role of Uterine Fibroids in Endometrial Function. Reprod Sci 25, 635–643 (2018).

51. Sinclair, D.C., Mastroyannis, A. & Taylor, H.S. Leiomyoma simultaneously impair endometrial BMP-2-mediated decidualization and anticoagulant expression through secretion of TGF-beta3. J Clin Endocrinol Metab 96, 412–421 (2011).

52. Wang, W., et al. Single-cell transcriptomic atlas of the human endometrium during the menstrual cycle. Nat Med 26, 1644–1653 (2020).

53. Jin, S., et al. Inference and analysis of cell-cell communication using CellChat. Nat Commun 12, 1088 (2021).

54. Paul, E.N., et al. Cysteine-rich intestinal protein 1 is a novel surface marker for human myometrial stem/progenitor cells. Commun Biol 6, 686 (2023).

55. Goad, J., et al. Single-cell sequencing reveals novel cellular heterogeneity in uterine leiomyomas. Hum Reprod 37, 2334–2349 (2022).

56. Arici, A. & Sozen, I. Transforming growth factor-beta3 is expressed at high levels in leiomyoma where it stimulates fibronectin expression and cell proliferation. Fertil Steril 73, 1006–1011 (2000).

57. Lee, B.S. & Nowak, R.A. Human leiomyoma smooth muscle cells show increased expression of transforming growth factor-beta 3 (TGF beta 3) and altered responses to the antiproliferative effects of TGF beta. J Clin Endocrinol Metab 86, 913–920 (2001).

58. Hartung, I.V., et al. Modular Assembly of Allosteric MEK Inhibitor Structural Elements Unravels Potency and Feedback-Modulation Handles. ChemMedChem 10, 2004–2013 (2015).

59. 59. Wyman, D., et al. A technology-agnostic long-read analysis pipeline for transcriptome discovery and quantification. bioRxiv (2020).

60. Reese, F. & Mortazavi, A. Swan: a library for the analysis and visualization of long-read transcriptomes. Bioinformatics 37, 1322–1323 (2021).

61. Geuens, T., Bouhy, D. & Timmerman, V. The hnRNP family: insights into their role in health and disease. Hum Genet 135, 851–867 (2016).

62. Thibault, P.A., et al. hnRNP A/B Proteins: An Encyclopedic Assessment of Their Roles in Homeostasis and Disease. Biology (Basel) 10(2021).

63. Alarcon, C.R., et al. HNRNPA2B1 Is a Mediator of m(6)A-Dependent Nuclear RNA Processing Events. Cell 162, 1299–1308 (2015).

64. Pankov, R. & Yamada, K.M. Fibronectin at a glance. J Cell Sci 115, 3861–3863 (2002).

65. Dalton, C.J. & Lemmon, C.A. Fibronectin: Molecular Structure, Fibrillar Structure and Mechanochemical Signaling. Cells 10(2021).

66. Spada, S., Tocci, A., Di Modugno, F. & Nistico, P. Fibronectin as a multiregulatory molecule crucial in tumor matrisome: from structural and functional features to clinical practice in oncology. J Exp Clin Cancer Res 40, 102 (2021).

67. Dorafshan, S., et al. Periostin: biology and function in cancer. Cancer Cell Int 22, 315 (2022).

68. Menning, A., et al. Granulocytes and vascularization regulate uterine bleeding and tissue remodeling in a mouse menstruation model. PLoS One 7, e41800 (2012).

69. Islam, M.S., Ciavattini, A., Petraglia, F., Castellucci, M. & Ciarmela, P. Extracellular matrix in uterine leiomyoma pathogenesis: a potential target for future therapeutics. Hum Reprod Update 24, 59–85 (2018).

70. Navarro, A., Bariani, M.V., Yang, Q. & Al-Hendy, A. Understanding the Impact of Uterine Fibroids on Human Endometrium Function. Front Cell Dev Biol 9, 633180 (2021).

71. Hynes, R.O. & Naba, A. Overview of the matrisome--an inventory of extracellular matrix constituents and functions. Cold Spring Harb Perspect Biol 4, a004903 (2012).

72. Wegienka, G. Are uterine leiomyoma a consequence of a chronically inflammatory immune system? Med Hypotheses 79, 226–231 (2012).

73. Fletcher, N.M., et al. Uterine fibroids are characterized by an impaired antioxidant cellular system: potential role of hypoxia in the pathophysiology of uterine fibroids. J Assist Reprod Genet 30, 969–974 (2013).

74. Donnez, J. & Dolmans, M.M. Uterine fibroid management: from the present to the future. Hum Reprod Update 22, 665–686 (2016).

75. Okada, H., Tsuzuki, T. & Murata, H. Decidualization of the human endometrium. Reprod Med Biol 17, 220–227 (2018).

76. Stewart, E.A., Friedman, A.J., Peck, K. & Nowak, R.A. Relative overexpression of collagen type I and collagen type III messenger ribonucleic acids by uterine leiomyomas during the proliferative phase of the menstrual cycle. J Clin Endocrinol Metab 79, 900–906 (1994).

77. Behera, M.A., et al. Thrombospondin-1 and thrombospondin-2 mRNA and TSP-1 and TSP-2 protein expression in uterine fibroids and correlation to the genes COL1A1 and COL3A1 and to the collagen cross-link hydroxyproline. Reprod Sci 14, 63–76 (2007).

78. Moore, S.C., Gierach, G.L., Schatzkin, A. & Matthews, C.E. Physical activity, sedentary behaviours, and the prevention of endometrial cancer. Br J Cancer 103, 933–938 (2010).

79. Iwahashi, M., et al. Immunohistochemical analysis of collagen expression in uterine leiomyomata during the menstrual cycle. Exp Ther Med 2, 287–290 (2011).

80. Malik, M., Norian, J., McCarthy-Keith, D., Britten, J. & Catherino, W.H. Why leiomyomas are called fibroids: the central role of extracellular matrix in symptomatic women. Semin Reprod Med 28, 169–179 (2010).

81. Bogusiewicz, M., et al. Expression of matricellular proteins in human uterine leiomyomas and normal myometrium. Histol Histopathol 27, 1495–1502 (2012).

82. Patel, A., Malik, M., Britten, J., Cox, J. & Catherino, W.H. Mifepristone inhibits extracellular matrix formation in uterine leiomyoma. Fertil Steril 105, 1102–1110 (2016).

83. Shen, X., Yang, Z., Feng, S. & Li, Y. Identification of uterine leiomyosarcoma-associated hub genes and immune cell infiltration pattern using weighted co-expression network analysis and CIBERSORT algorithm. World J Surg Oncol 19, 223 (2021).

84. Courtoy, G.E., et al. Gene expression changes in uterine myomas in response to ulipristal acetate treatment. Reprod Biomed Online 37, 224–233 (2018).

85. Malik, M., Britten, J., Borahay, M., Segars, J. & Catherino, W.H. Simvastatin, at clinically relevant concentrations, affects human uterine leiomyoma growth and extracellular matrix production. Fertil Steril 110, 1398–1407 e1391 (2018).

86. Black, D.L. Mechanisms of alternative pre-messenger RNA splicing. Annu Rev Biochem 72, 291–336 (2003).

87. Zhang, Y., Qian, J., Gu, C. & Yang, Y. Alternative splicing and cancer: a systematic review. Signal Transduct Target Ther 6, 78 (2021).

88. Cheng, Z., Shang, Y., Gao, S. & Zhang, T. Overexpression of U1 snRNA induces decrease of U1 spliceosome function associated with Alzheimer’s disease. J Neurogenet 31, 337–343 (2017).

89. Takayama, K.I., et al. Dysregulation of spliceosome gene expression in advanced prostate cancer by RNA-binding protein PSF. Proc Natl Acad Sci U S A 114, 10461–10466 (2017).

90. Zhao, Y. & Young, S.L. TGF-beta regulates expression of tenascin alternative-splicing isoforms in fetal rat lung. Am J Physiol 268, L173–180 (1995).

91. Tripathi, V. & Zhang, Y.E. Redirecting RNA splicing by SMAD3 turns TGF-beta into a tumor promoter. Mol Cell Oncol 4, e1265699 (2017).

92. Weg-Remers, S., Ponta, H., Herrlich, P. & Konig, H. Regulation of alternative pre-mRNA splicing by the ERK MAP-kinase pathway. EMBO J 20, 4194–4203 (2001).

93. Pelisch, F., Blaustein, M., Kornblihtt, A.R. & Srebrow, A. Cross-talk between signaling pathways regulates alternative splicing: a novel role for JNK. J Biol Chem 280, 25461–25469 (2005).

94. Goncalves, V., Matos, P. & Jordan, P. Antagonistic SR proteins regulate alternative splicing of tumor-related Rac1b downstream of the PI3-kinase and Wnt pathways. Hum Mol Genet 18, 3696–3707 (2009).

95. Chang, J.W., et al. mTOR-regulated U2af1 tandem exon splicing specifies transcriptome features for translational control. Nucleic Acids Res 47, 10373–10387 (2019).

96. Lee, F.F., et al. NF-kappaB mediates lipopolysaccharide-induced alternative pre-mRNA splicing of MyD88 in mouse macrophages. J Biol Chem 295, 6236–6248 (2020).

97. Chu, W.K., Hung, L.M., Hou, C.W. & Chen, J.K. PKC Regulates YAP Expression through Alternative Splicing of YAP 3’UTR Pre-mRNA by hnRNP F. Int J Mol Sci 22(2021).

98. Iborra, A., Mayorga, M., Llobet, N. & Martinez, P. Expression of complement regulatory proteins [membrane cofactor protein (CD46), decay accelerating factor (CD55), and protectin (CD59)] in endometrial stressed cells. Cell Immunol 223, 46–51 (2003).

99. Nogawa Fonzar-Marana, R.R., et al. Expression of complement system regulatory molecules in the endometrium of normal ovulatory and hyperstimulated women correlate with menstrual cycle phase. Fertil Steril 86, 758–761 (2006).

100. Hiroi, H., et al. Expression and regulation of periostin/OSF-2 gene in rat uterus and human endometrium. Endocr J 55, 183–189 (2008).

101. Xu, X., et al. Periostin Enhances Migration, Invasion, and Adhesion of Human Endometrial Stromal Cells Through Integrin-Linked Kinase 1/Akt Signaling Pathway. Reprod Sci 22, 1098–1106 (2015).

102. Cao, W., Mah, K., Carroll, R.S., Slayden, O.D. & Brenner, R.M. Progesterone withdrawal up-regulates fibronectin and integrins during menstruation and repair in the rhesus macaque endometrium. Hum Reprod 22, 3223–3231 (2007).

103. Bilalis, D.A., Klentzeris, L.D. & Fleming, S. Immunohistochemical localization of extracellular matrix proteins in luteal phase endometrium of fertile and infertile patients. Hum Reprod 11, 2713–2718 (1996).

104. Bordeleau, F., et al. Tissue stiffness regulates serine/arginine-rich protein-mediated splicing of the extra domain B-fibronectin isoform in tumors. Proc Natl Acad Sci U S A 112, 8314–8319 (2015).

105. Danecek, P., et al. Twelve years of SAMtools and BCFtools. Gigascience 10(2021).

106. Cingolani, P., et al. A program for annotating and predicting the effects of single nucleotide polymorphisms, SnpEff: SNPs in the genome of Drosophila melanogaster strain w1118; iso-2; iso-3. Fly (Austin) 6, 80–92 (2012).

107. Cameron, D.L., et al. GRIDSS2: comprehensive characterisation of somatic structural variation using single breakend variants and structural variant phasing. Genome Biol 22, 202 (2021).

108. Love, M.I., Huber, W. & Anders, S. Moderated estimation of fold change and dispersion for RNA-seq data with DESeq2. Genome Biol 15, 550 (2014).

109. Yu, G., Wang, L.G., Han, Y. & He, Q.Y. clusterProfiler: an R package for comparing biological themes among gene clusters. OMICS 16, 284–287 (2012).

110. Wu, T., et al. clusterProfiler 4.0: A universal enrichment tool for interpreting omics data. Innovation (Camb) 2, 100141 (2021).

111. Soneson, C., Love, M.I. & Robinson, M.D. Differential analyses for RNA-seq: transcript-level estimates improve gene-level inferences. F1000Res 4, 1521 (2015).

112. Nowicka, M. & Robinson, M.D. DRIMSeq: a Dirichlet-multinomial framework for multivariate count outcomes in genomics. F1000Res 5, 1356 (2016).

113. Anders, S., Reyes, A. & Huber, W. Detecting differential usage of exons from RNA-seq data. Genome Res 22, 2008–2017 (2012).

114. Kuleshov, M.V., et al. Enrichr: a comprehensive gene set enrichment analysis web server 2016 update. Nucleic Acids Res 44, W90–97 (2016).

115. Amezquita, R.A., et al. Orchestrating single-cell analysis with Bioconductor. Nat Methods 17, 137–145 (2020).

116. Hao, Y., et al. Integrated analysis of multimodal single-cell data. Cell 184, 3573–3587 e3529 (2021).

117. Germain, P.L., Lun, A., Garcia Meixide, C., Macnair, W. & Robinson, M.D. Doublet identification in single-cell sequencing data using scDblFinder. F1000Res 10, 979 (2021).

118. Korsunsky, I., et al. Fast, sensitive and accurate integration of single-cell data with Harmony. Nat Methods 16, 1289–1296 (2019).

119. Li, H. Minimap2: pairwise alignment for nucleotide sequences. Bioinformatics 34, 3094–3100 (2018).

120. Cohen, P.E. & Milligan, S.R. Silastic implants for delivery of oestradiol to mice. J Reprod Fertil 99, 219–223 (1993).

121. Hallberg, L. & Nilsson, L. Determination of Menstrual Blood Loss. Scand J Clin Lab Invest 16, 244–248 (1964).

